# Intensity Modulation of Trichromatic Split Fluorescent Proteins for Live Cell Mapping

**DOI:** 10.1101/2025.05.13.653616

**Authors:** Mamoru Ishii, Tomoaki Kinjo, Yohei Kondo, Kenta Terai, Kazuhiro Aoki, Brian Kuhlman, Michiyuki Matsuda

**Affiliations:** Graduate School of Biostudies, Kyoto University, Sakyo-ku, Kyoto 606-8501, Japan; Institute of Industrial Science, The University of Tokyo, Tokyo, Japan; Department of Biochemistry and Biophysics, University of North Carolina School of Medicine, Chapel Hill, NC, USA; Center for One Medicine Innovative Translational Research (COMIT), Nagoya University, Nagoya, Aichi, Japan; Graduate School of Medicine, Nagoya University, Tsurumai-cho, Nagoya, Aichi 466-8550, Japan; Graduate School of Medicine, Tokushima University, Shinkura-cho, Tokushima 770-8501, Japan; Lineberger Comprehensive Cancer Center, University of North Carolina at Chapel Hill, Chapel Hill, NC, USA; Affiliate Graduate School, Graduate School of Medicine, Kyoto University, Sakyo-ku, Kyoto 606-8501, Japan; Integrated Graduate School of Medicine, Engineering, and Agricultural Sciences, University of Yamanashi, Chuo-shi, Yamanashi 409-3898 Japan

**Keywords:** Split fluorescent proteins, Multiplexed spectral labeling, Cell labeling, Fluorescence microscopy, Protein tagging

## Abstract

Multiplexed live cell-labeling technologies are essential tools for investigating complex biological systems, yet existing approaches face limitations in achieving reliable discrimination of multiple cell populations. Here, we present color and intensity modulation with split fluorescent proteins for cell labeling (Caterpie), a live cell labeling system that enables high-fidelity identification of multiple cell populations through rational design of split fluorescent proteins. Drawing inspiration from human trichromatic vision, we engineered optimized split fluorescent protein variants—split CFP2, split mNG3A, and split sfCherry3C—and developed a comprehensive library of the 11th β-strand (FP_11_) tags comprising variable numbers of tandem repeats. Structure-guided protein engineering enhanced the performance of split mNG3A and split sfCherry3C through targeted modifications of their C-terminal sequences and binding interfaces, respectively. By systematically varying the composition and arrangement of FP_11_ repeats, we selected 20 distinct FP_11_ tags that enable robust cell population discrimination with 97% accuracy using conventional fluorescence microscopy. In these FP_11_ tag-expressing cells, we expressed multiple EGF ligands and their receptors and cultured them in bulk to examine the effects of EGF signaling on chemotaxis within the cell population. The results showed that chemotaxis changes according to the intensity of EGFR signaling. The Caterpie system holds significant potential for applications requiring precise identification and tracking of multiple cell populations in complex biological systems.

## Introduction

Multiplexed spectral labeling techniques represent essential tools across diverse biological disciplines, including neuroscience, developmental biology, and immunology. These techniques enable critical applications such as discrimination of adjacent cells, specific cell populations labeling, and lineage tracing. The Brainbow system exemplifies this approach, utilizing multiple gene cassettes designed to stochastically express one of three fluorescent proteins (FPs), thereby generating distinct spectral signatures (*1*). While this system has proven particularly valuable for morphological analyses including connectomics, it presents limitations for specific cell labeling and lineage tracing due to potential variations in the relative FP expression levels across cellular generations. The development of Bitbow addressed certain limitations of earlier systems by enabling the generation of up to 31 (2^5^-1) distinct color variations through random toggling of five FP genes within a single gene cassette (*2*). However, this approach faces challenges due to biased gene selection partners, resulting in fewer achievable color discrimination than theoretically predicted. These limitations underscore the advantages of rationally designed color palettes. However, conventional approaches to expanding color diversity through serial concatenation of multiple FPs face practical constraints due to increasing gene sizes. Split fluorescent proteins emerge as an elegant solution to these technical challenges.

Split fluorescent proteins (split FPs) are generated through strategic dissection of β-barrel fluorescent proteins (FPs) into two components: the 11th β-strand (FP_11_) and a complementary segment comprising the first ten β-strands (FP_1-10_) (*3*). While neither fragment exhibits fluorescence independently, their co-expression facilitates spontaneous reassembly, enabling chromophore maturation and subsequent fluorescence emission. This complementation system has been successfully employed for protein labeling applications, wherein target proteins are tagged with FP_11_ in cells expressing FP_1-10_ (*4*, *5*). A particularly advantageous feature of this system emerges under conditions of abundant FP_1-10_ expression: enhanced fluorescence intensity can be achieved by increasing the number of FP_11_ tags conjugated to the protein of interest, while maintaining relatively modest gene size requirements (*6*–*9*).

Drawing inspiration from human color perception, which discriminates diverse spectral signatures through the integration of three primary color intensities: blue, green, and red, we developed a novel multiplexed cellular labeling approach. This system exploits engineered arrays of FP_11_ fragments derived from cyan, green, and red split fluorescent proteins, each designed to complement with their cognate FP_1-10_ fragments. This method, named color and intensity modulation with split fluorescent proteins for cell labeling (Caterpie), achieves robust discrimination of 20 distinct cell populations with 97% accuracy using conventional epi-fluorescence microscopy.

## Results

### Split fluorescent proteins for the trichromatic color palette

We aimed to establish a trichromatic color palette through the systematic engineering of tandem FP_11_ repeats derived from cyan, green, and red split fluorescent proteins, which undergo fluorescence complementation when co-expressed with their cognate FP_1-10_ fragments (Fig. 1A). For this, we sought to select split fluorescent proteins (split FPs) that achieve brightness comparable to their full-length FP (FP_Full-length_) and exhibit fluorescence signal amplification through FP_11_ repeats. A critical consideration in designing this system is the requirement for sequence divergence among the trichromatic split FPs to prevent cross-complementation between fragments from different fluorescent protein pairs (*10*). To address this constraint, we evaluated split FPs derived from distinct evolutionary lineages: sfGFP, mNG2, sfCherry2, and mRuby4 (Fig. 1B). To quantitatively assess the relative brightness of FP_Full-length_, FP_11_(x1), and FP_11_(x4) of reported split FPs, we developed a dual-reporter system. We engineered expression constructs encoding Histone H2B-tagged FP_Full-length_, FP_11_(x1), or FP_11_(x4) fused to EBFP2-nls (nuclear localization signal) through a self-cleaving P2A site, enabling normalization of expression levels through nuclear EBFP2 fluorescence. These constructs were co-transfected into HeLa cells with a secondary plasmid encoding FP_1-10_ linked to iRFP670 via an internal ribosome entry site (IRES) at a DNA ratio of 1:4 (Fig. 1C).

**Figure 1.**
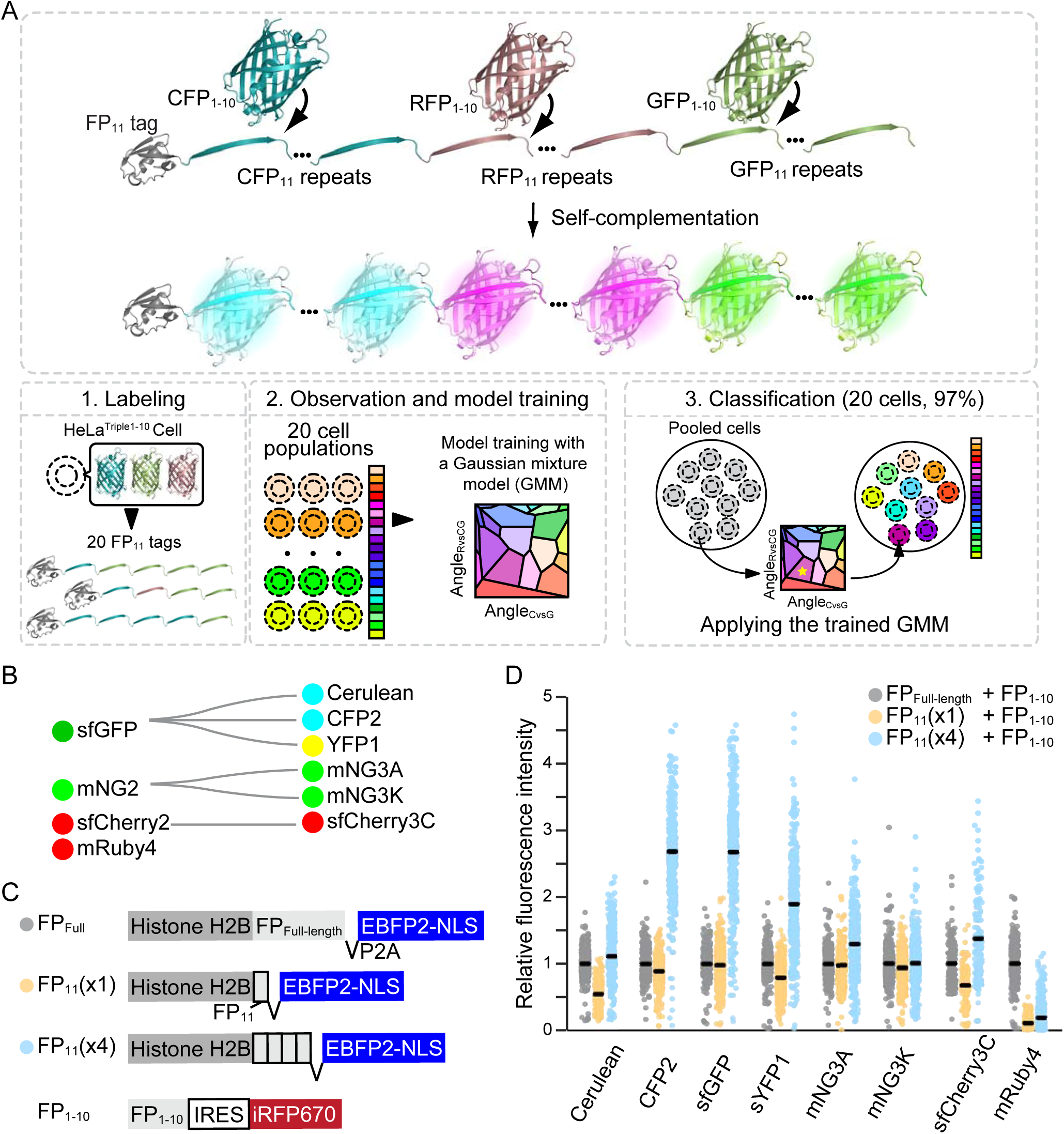
Caterpie: A cell identification system based on split fluorescent protein arrays. **(A)** Schematic overview of the Caterpie approach, illustrating cell identification through modular arrays of split fluorescent proteins. **(B)** Evolutionary relationships among split fluorescent proteins utilized in this study, presented as a phylogenetic tree. **(C)** Molecular architecture of expression constructs: Histone H2B fusions containing either full-length FP (FP_Full-length_), single FP_11_ fragment, or tetrameric FP_11_ arrays, shown alongside the complementary FP_1-10_ fragment. **(D)** Quantitative analysis of relative fluorescence intensity in HeLa cells expressing FP_Full-length_, FP_11_(x1) or FP_11_(x4), with FP_1-10_ fragments (1:4 transfection ratio). Data presented as bee swarm plots with median values indicated by black lines (n > 130 cells per condition). Measurements obtained through confocal microscopy.

After 48 hours, cells were observed under a confocal microscope, and split FP fluorescence intensities were quantified in iRFP670 positive cells (Fig. 1D). For each fluorescent protein variant, relative fluorescence intensity values were normalized to the full-length FP. In most cases, (x1) shows decreased fluorescence brightness compared to full-length, but sfGFP, mNG3A, and mNG3K were almost equivalent fluorescence brightness. Comparative analysis of (x1) versus (x4) constructs revealed that all sfGFP-derived split FPs exhibited more than two-fold enhancement in median brightness, with CFP2 demonstrating the most substantial increase of more than three-fold. Consistent with previous findings (*11*), split mNG3A demonstrated superior amplification from (x1) to (x4) compared to split mNG3K. Split sfCherry3C_11_(x1) exhibited diminished fluorescence intensity relative to sfCherry3C_Full-length_, attributable to reduced association efficiency, corroborating earlier observations (*12*). The truncated mRuby4_11_ showed markedly decreased fluorescence intensity, highlighting the critical role of its C-terminal unstructured polypeptide chain (*10*). These preliminary findings indicated that while the combination of CFP2, mNG3A, and sfCherry3C provides a promising foundation for the trichromatic system, although further engineering optimization of mNG3A and sfCherry3C is necessary to achieve optimal performance across all spectral channels.

### Fluorescence intensity-based cell classification using tandem CFP2 **β**-strand repeats

Before the engineering of mNG3A and sfCherry3C, we established a platform for cell identification through fluorescence intensity modulation using CFP2_11_ repeats. To this end, we constructed plasmids encoding repeated sequences of CFP2_11_ (Fig. 2A). Initially, we engineered a plasmid encoding a Histone H2B-CFP2_11_(x1) fusion construct, in which the CFP2_11_(x1) sequence was flanked by a BglII restriction site at its 5’ terminus and BamHI and NotI restriction sites at its 3’ terminus. Through sequential ligation of BglII/NotI-digested fragments (insert) with BamHI/NotI-digested fragments (vector), we successfully amplified CFP2_11_ to create (x2), (x4), and (x8) variants. Based on previous research (*6*), we incorporated GGSGG linker sequences between CFP2_11_ fragments. At the BglII-BamHI junction, the six nucleotides encode glycine and serine residues, thereby serving as an integral part of the linker sequence. The CFP2_11_ repeats [(x1), (x2), (x4), or (x8)] fused with Histone H2B were introduced into HeLa cells stably expressing CFP2_1-10_ (HeLa^CFP2^ ^1-10^) (Fig. 2B). As a control, we expressed Histone H2B-tagged CFP2_Full-length_ in the same HeLa^CFP2^ ^1-10^ cell line. When normalized to nuclear mCherry fluorescence, the CFP2 fluorescence intensity increased across consecutive constructs with a factor of approximately 1.7-fold, which was slightly lower than the theoretically expected 2-fold increment (Fig. 2C).

**Figure 2.**
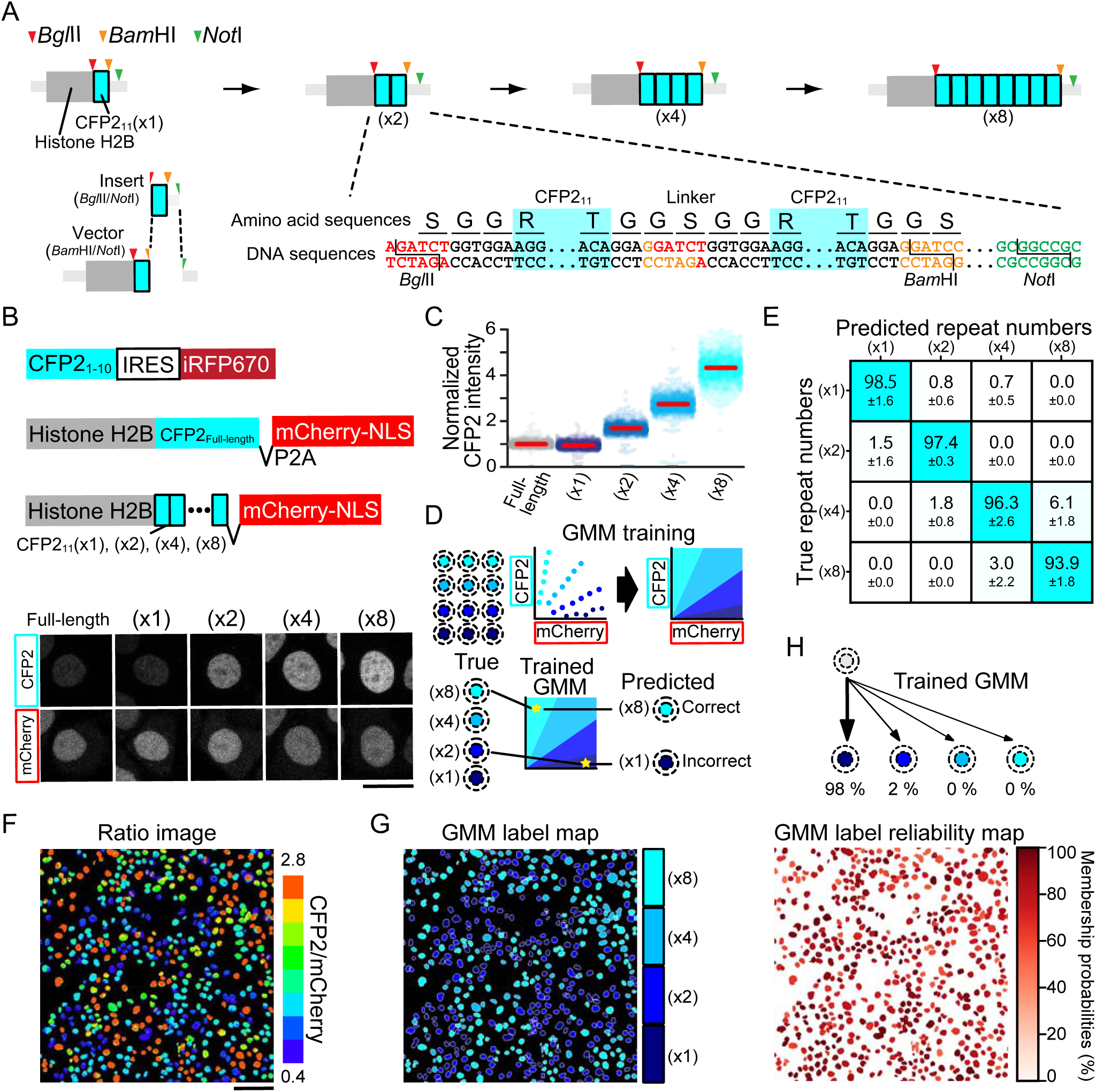
Signal amplification and cell classification using split CFP2 arrays. **(A)** Molecular strategy for generating tandem CFP2_11_ repeats, including detailed amino acid and nucleotide sequences at the CFP2_11_(x1) insert-vector junction. **(B)** Construct architecture and expression analysis: Upper panel shows schematics CFP2_1-10_ and Histone H2B fusions containing either full-length CFP2 (CFP2_Full-length_) or varying copy numbers of CFP2_11_ [CFP2_11_(x1), (x2), (x4), and (x8)]. Lower panel presents representative confocal micrographs of HeLa cells stably co-expressing CFP2_1-10_ with either CFP2_Full-length_ or CFP2_11_ variants. Scale bar, 20□µm. **(C)** Quantitative analysis of normalized CFP2 fluorescence in HeLa cells expressing Histone H2B-tagged CFP2_11_ [(x1), (x2), (x4), or (x8)]. Data presented as bee swarm plots with median values (red lines); >1300 cells analyzed across three independent experiments. **(D)** Workflow schematic for Gaussian mixture model (GMM) and implementation. **(E)** Classification performance matrix showing prediction accuracy against true labels (rows). Data represented as mean ± SD of prediction accuracy from three independent experiments, with color intensity indicating classification accuracy. **(F)** Representative ratio image of pooled cells expressing different CFP2_11_ copy number variants [(x1), (x2), (x4), or (x8)]; scale bar: 100□µm. **(G)** Cell classification map derived from panel (F), showing GMM-based assignment of individual cells to specific copy number variants [(x1), (x2), (x4), or (x8)]. **(H)** Visualization of GMM classification confidence through membership probability mapping of cells shown in panel (G).

To identify cells expressing different numbers of CFP2_11_ repeats, we implemented a Gaussian mixture model (GMM) for cell classification (Fig. 2D). The fluorescence intensities of mCherry and CFP2 were transformed into polar coordinates, where the Angle_mCherry vs CFP2_ (ranging from 0° to 90°) represented the CFP2/mCherry intensity ratio. The angular data were divided into training and test datasets. We trained a GMM on the training dataset to establish classification parameters, determining the decision boundary of the Angle_mCherry vs CFP2_. The model performance was validated using the test dataset, and the predicted repeat numbers were compared with the actual repeat numbers. The model achieved an overall accuracy of 96% (Fig. 2E). Among all constructs, CFP2_11_(x8) exhibited the lowest accuracy (93.9%), with 6.1% of (x8)-expressing cells being misclassified as (x4).

To validate the system’s discriminative capacity in a heterogeneous context, we analyzed mixed populations of cells expressing four different CFP2_11_ repeat variants using fluorescence microscopy and generated CFP2/mCherry ratio images (Fig. 2F). The trained GMM demonstrated robust performance in classifying individual cells into (x1), (x2), (x4), or (x8) populations with 96% accuracy, which was validated through visual inspection (Fig. 2G).

Furthermore, the GMM provided membership probabilities for each cell, enabling quantitative assessment of classification reliability (Fig. 2H). These results established that four-level intensity modulation of a single fluorescent protein is sufficient for reliable population discrimination.

### Computational design of split sfCherry3C for enhanced **β**-barrel stability and split-strand complementation

To optimize split sfCherry3C toward fluorescence intensities close to those of the full-length protein, we modelled the sfCherry3C_1-10_/sfCherry3C_11_ complex with ColabFold (*13*) (based on AlphaFold-Multimer) (*14*), and used the structure model for subsequent design with Rosetta modeling suite (*15*). Because both global β-barrel stability and β_1-10_/β_11_ interface affinity are crucial for fluorescence of split FPs, we pursued two design strategies in parallel (*16*). First, to enhance global β-barrel stability, we employed a Rosetta script adapted from the previously described “Point-Mutation” (PM) workflow (*16*) and performed *in-silico* site-saturation mutagenesis (SSM) across all 224 residues, ranking each variant by the change in calculated energy (ΔE) calculated from the Rosetta *total_score* term (fig. S1). Next, to refine the β_1-10_/β_11_ interface, we repurposed the PM script for interface design; with the revised script, we performed SSM at every β_11_ position and calculated the binding energy of each single mutant from *dG_separated* score obtained with InterfaceAnalyzer protocol (Fig. 3A). Each single-mutant model was subsequently processed by the “Mutation Cluster (MC)” (*16*) workflow modified for interface design, in which FastDesign protocol with InterfaceDesign relax script stochastically samples sequence and rotamer space at residues within 5–7 Å of the “seed mutation” through Monte-Carlo design/minimization cycles, yielding multi-residue mutants that are re-evaluated with InterfaceAnalyzer (Fig. 3B). After filtering by the scores such as *total_score*, *dG_separated*, and mutation number, we conducted visual inspection to retain the variants that reinforce interface contacts while excluding ones that disrupted the chromophore-forming triad M67–Y68–G69 or introduced additional aromatic residues into β_11_ which could promote aggregation of multimeric β_11_ repeat. Following these criteria, we selected twenty-two candidate designs for subsequent in-cell validation.

**Figure 3.**
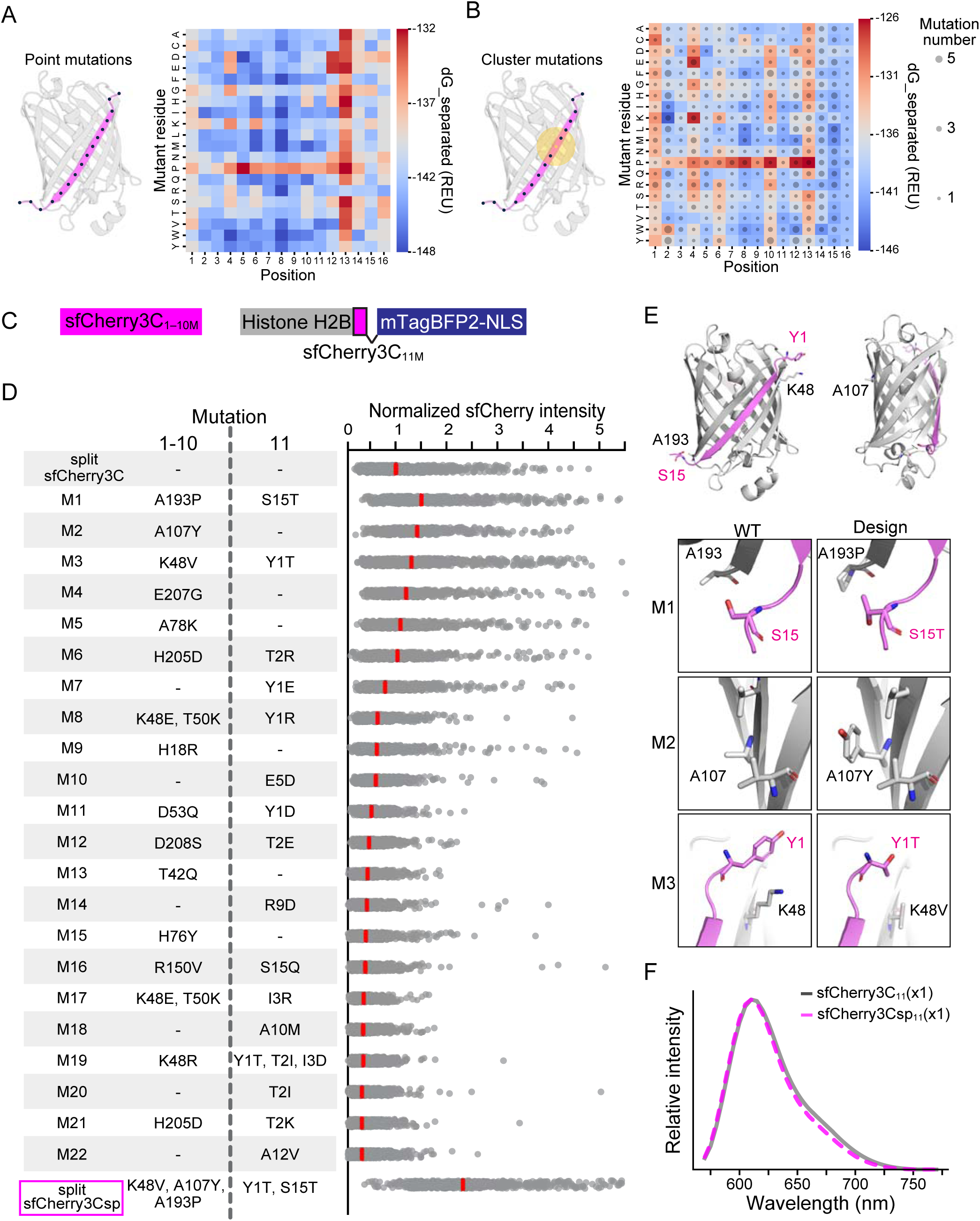
Computational design of split sfCherry3C. (A) In-silico site-saturation mutagenesis of sfCherry3C□□. Left: schematic of the β□□ residues targeted for saturation mutagenesis (black spheres). Right: heatmap of mean dG_separated values in Rosetta energy unit (REU) across three independent models for each amino acid substitution at each position. (B) Cluster-mutation design of sfCherry3C□□. Left: schematic showing a representative seed point mutation (yellow dot) and the shell of neighboring residues (yellow spheres) targeted for sequential cluster mutagenesis. Right: heatmap of the mean dG_separated across ten cluster-mutated models derived from each seed mutation at every β□□ site; dot size indicates the median number of cluster-mutated residues across these ten models. (C) Schematics of mutant variants of sfCherry3C_1-10_ (sfCherry3C_1-10M_) and Histone H2B-tagged mutant variant of sfCherry3C_11_ (sfCherry3C_11M_). (D) List of mutant variants of sfCherry3C_1-10M_ and sfCherry3C_11M_. Bee swarm plot showing normalized sfCherry3C fluorescence intensity of HeLa cells stably co-expressing Histone H2B-tagged sfCherry3C_11M_(x1) with sfCherry3C_1-10M_. Red lines represent the median. (E) Rosetta models of split sfCherry3Csp. The overall β-barrel is shown as a cartoon colored by fragment (gray, β□–□□; magenta, β□□). Each inset zooms on an engineered site, showing the parental conformation on the left and the corresponding design on the right, with key residues rendered as sticks representation. All models were visualized in PyMOL. (F) Emission spectra of HeLa cells expressing split sfCherry3C or split sfCherry3Csp with 546 nm laser excitation.

Upon stable co-expression of β_1-10_ and β_11_ in HeLa cells, three variants (M1, M2, M3) outperformed the parental split sfCherry3C in fluorescent intensity (Fig. 3C, D). We then combined the mutations from the three top performing variants to generate an optimized variant, named split sfCherry3Csp, which achieved a 2.5-fold enhancement in fluorescent intensity compared to the parental split sfCherry3C. The M1 and M3 mutations locate at the β_1-10_/β_11_ interface, where the models suggest the improved association of β_1-10_ and β_11_ through the formation of reinforced side-chain contacts. The M2 mutation lies on the opposite face of the interface, implying an effect on β-barrel stability (Fig. 3E). The fluorescence spectra of split sfCherry3Csp were indistinguishable to those of the parental split sfCherry3C, indicating that the incorporated mutations enhanced the β-barrel stability and β_1-10_/β_11_ association without affecting the chromophore maturation (Fig. 3F).

### Enhancement of complementation efficiency through structure-guided engineering of split mNG3A

As shown in Fig. 1D, split mNG3A demonstrated only modest signal amplification, with a 1.3-fold increase in fluorescence intensity from (x1) to (x4) variants. To gain insight into the structural factors underlying this limitation, we modelled the octameric mNG3A_11_ peptide using ColabFold (*13*) (based on AlphaFold2) (*17*). The predicted structures suggested that mNG3A_11_ repeat adopts a loosely helical conformation, with hydrophobic residues (F5, W8, F12, M15, M16) converging to form a continuous hydrophobic interaction, despite only moderate model confidence. These features implied that multimeric mNG3A_11_ may self-aggregate via its hydrophobic surfaces (Fig. 4A) (*17*). Additionally, inspection of the parental mNG crystal structure (PDB: 5LTP) and the structure model of split mNG3A suggested that the C-terminal residues D14–M15–M16 contribute minimally to the β-barrel fold (Fig. 4B). We therefore reasoned that removing these C-terminal residues could mitigate the aggregation propensity without perturbing β-barrel and chromophore maturation. Guided by these insights, we implemented three C-terminal modification approaches: (1) deletion of the C-terminal D14-M15-M16, (2) addition of one or two aspartic acid residue to preserve charge that could electrostatically repel neighboring fragments, and (3) substitution of F12 with tyrosine to introduce polarity (Fig. 4C). To evaluate these modifications, we expressed 8 repeats of the mNG3A_11_ fragments fused to Histone H2B in HeLa cells expressing mNG3A_1-10_ (Fig. 4D). Nuclear mNG3A fluorescence was quantified through microscopy, with expression levels normalized to nuclear mTagBFP2 fluorescence intensity (Fig. 4E). Among the modifications tested, we chose the mNG3Asp_11_ variant for further analysis because of its highest fluorescence intensity, which was 9.1-fold brighter than the parental mNG3A_11_. The fluorescence spectra obtained using the mNG3Asp_11_ fragment were identical to those observed with the mNG3A_11_ fragment (Fig. 4F).

**Figure 4.**
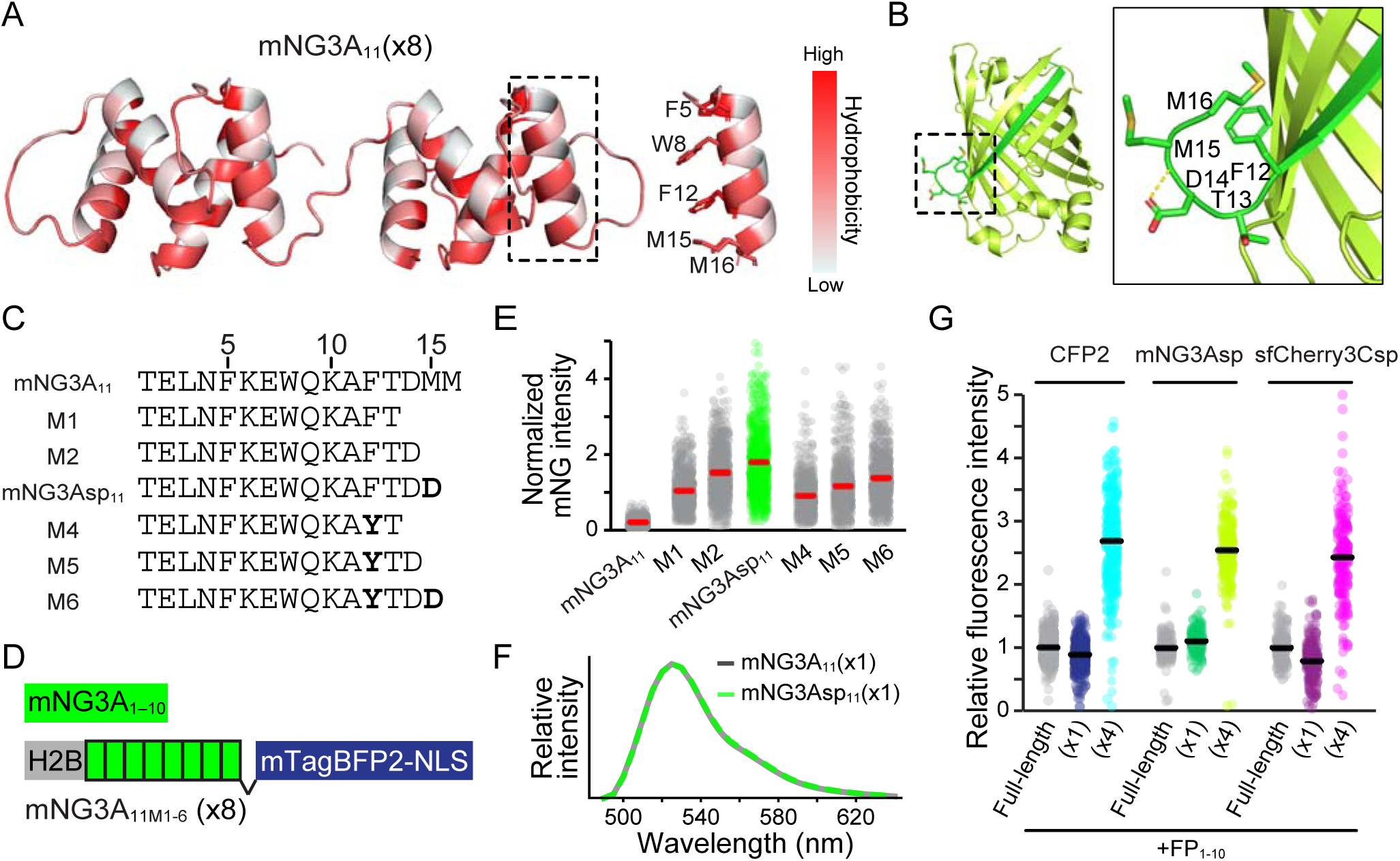
Optimization of split mNG3A through structure-guided C-terminal engineering. **(A)** Predicted structural model of native mNG3A11(x8) generated using ColabFold (AlphaFold2) and visualized with PyMOL. Colors represent hydrophobicity according to the Eisenberg hydrophobicity scale. **(B)** Predicted structural model of parental split mNG3A generated using ColabFold (AlphaFold Multimer) and visualized with PyMOL. Color-coded domains: mNG3A_1-10_ (green) and mNG3A_11_ (light green). **(C)** Comparative sequence analysis of original mNG3A_11_ and engineered mutant variants (mNG3A_11M_). **(D)** Schematic representation of expression constructs: mNG3A_1-10_ and Histone H2B fusion with the mNG3A_11M_(x8) array. Quantitative analysis of normalized mNG3A fluorescence in HeLa cells expressing Histone H2B-tagged mNG3A_11_ [(x1), (x2), (x4), or (x8)]. Data presented as bee swarm plots with median values (red lines). **(E)** Quantitative analysis of normalized mNG3A fluorescence intensity in HeLa cells stably co-expressing Histone H2B-tagged mNG3A_11M_ (x8) with mNG3A_1-10_. Median values indicated by red lines. Values normalized to set the median value of mNG3A_11M1_(x8) to 1. **(F)** Emission spectra of HeLa cells expressing split mNG3A or split mNG3Asp with 470 nm laser excitation. **(G)** Comparative analysis of relative fluorescence intensity of HeLa cells expressing FP_Full-length_, FP_11_(x1) or FP_11_(x4), with cognate FP_1-10_ fragments (1:4 transfection ratio). Data presented as bee swarm plots with median values (red lines); n > 130 cells per condition. Measurements obtained through confocal microscopy.

Performance evaluation of split mNG3Asp and split sfCherry3Csp was conducted in parallel with split CFP2, following our previously established experimental framework (Fig. 4G). Quantitative analysis revealed that mNG3Asp_11_(x4) achieved a 2.3-fold enhancement in fluorescence intensity compared to its monomeric counterpart. The engineered sfCherry3Csp demonstrated robust performance, with its monomeric variant exhibiting 80% of the fluorescence intensity of the full-length protein, while the tetrameric construct showed a 3-fold signal amplification relative to the monomer. These significant improvements in signal amplification efficiency indicate that both engineered variants, split mNG3Asp and split sfCherry3Csp, now achieve performance metrics comparable to split CFP2, providing the basis for trichromatic imaging.

### Optimization of the fusion partner for the FP_11_ tag

Prior to establishing the FP_11_ tag library, we aimed to reduce construct size by identifying alternatives to the relatively large Histone H2B fusion tag. We evaluated multiple fusion protein configurations, comparing Histone H2B (126 amino acids) with two smaller alternatives: GB1 (56 amino acids) (*18*) and ΔSUMOstar (74 amino acids), a truncated version of SUMOstar (*19*), as well as constructs lacking fusion tags (fig. S2A). ΔSUMOstar was engineered by removing the unstructured polypeptide regions from both termini of SUMOstar; the resulting sequence is detailed in Figure S2B. In agreement with previous studies (*20*), constructs lacking fusion proteins demonstrated markedly reduced fluorescence intensity, an effect that persisted even in the octameric (x8) variant (fig. S2C). Comparative analysis of fusion partners revealed that ΔSUMOstar-tagged constructs achieved substantially higher fluorescence intensity than GB1-tagged variants in octameric configurations, demonstrating the importance of fusion partner selection for optimal signal amplification. The ΔSUMOstar-tagged constructs exhibited consistent signal enhancement, with fluorescence intensity increasing 1.6-fold for each doubling of CFP2_11_ repeats, ultimately enabling 93% discrimination accuracy between variants (fig. S2, D to F).

To optimize the modular arrangement of the three FP_11_ variants (CFP2_11_, mNG3Asp_11_, and sfCherry3Csp_11_), we conducted a systematic analysis of possible configurations. In initial experiments, we generated ΔSUMOstar fusion constructs containing six total repeats, wherein each FP_11_ fragment was represented in duplicate (fig. S3A). The complete set of six possible permutations was generated and stably expressed in HeLa cells constitutively expressing all three FP_1-10_ fragments. Quantitative analysis of relative fluorescence intensities was performed using EBFP2 fluorescence for normalization of expression levels (fig. S3B). Among all configurations evaluated, the CFP2_11_(x2)-sfCherry3Csp_11_(x2)-mNG3Asp_11_(x2) arrangement demonstrated marginally superior fluorescence intensity across all spectral channels, leading to its selection for subsequent studies.

### Selection of 20 optimal FP_11_ tags

Based on these preliminary findings, we established a comprehensive library of fusion constructs combining ΔSUMOstar with CFP2_11_, sfCherry3Csp_11_, and mNG3Asp_11_ fragments in this order (Fig. 5A). Each fragment was represented in varying copy numbers [(x0), (x1), (x2), (x4), or (x8)] for each. In the first screening, we limited the FP_11_ tag combinations to 96 variants, with the total number of repeats not exceeding (x12). The integrity of all FP_11_ repeat sequences in the 96 plasmid constructs was verified by Sanger sequencing, thereby establishing a comprehensive FP_11_ tag library.

**Figure 5.**
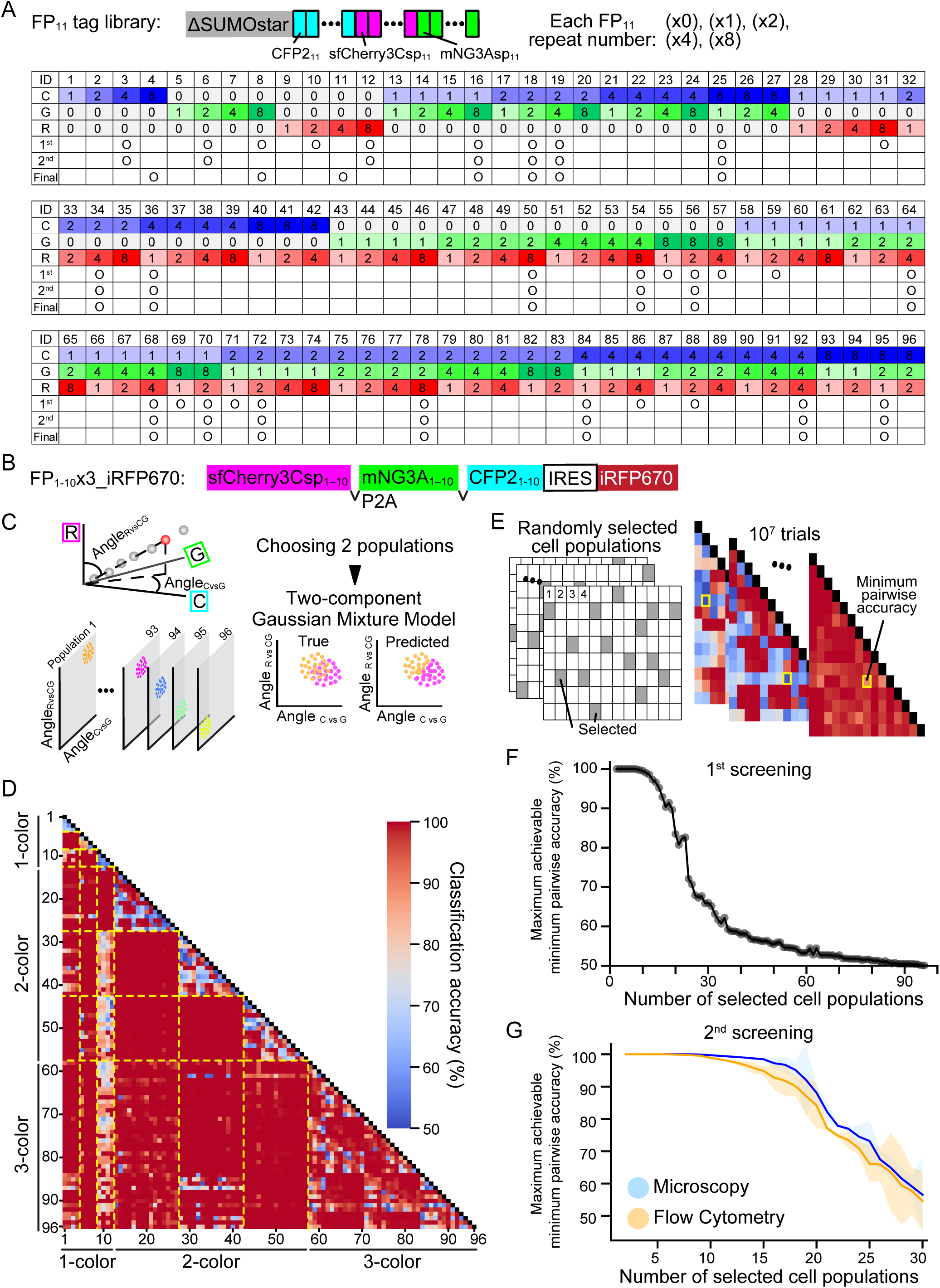
Systematic selection and validation of FP_11_ tags for Caterpie implementation. **(A)** Schematic representation of FP_11_ tags: ΔSUMO-tagged FP_11_ arrays comprising paired repeats of CFP2_11_, mNG3Asp_11_, and sfCherry3Csp_11_ [(x0), (x1), (x2), (x4), (x8) each]. Comprehensive catalog of the FP_11_ tag library comprising 96 distinct combinations, detailing copy numbers of CFP2_11_ (labeled as "C"), mNG3Asp_11_ (labeled as "G"), and sfCherry3Csp_11_ (labeled as "R") fragments. The marks indicate the FP_11_ tags selected during primary and secondary screening processes, and those finally chosen. **(B)** Schematic representation of expression constructs: FP_1-10_x3. **(C)** Analysis scheme for classification of cell populations expressing various FP_11_ tags. **(D)** Matrix analysis of pairwise classification accuracy across 96 cell populations expressing various FP_11_ tags, arranged by FP_11_ tag numbers shown in panel (A). (n > 230 cells per population). (**E)** Analysis scheme for determining the highest achievable minimum pairwise classification accuracy among randomly selected FP_11_-tagged subpopulations. **(F)** Graph showing the highest achievable minimum pairwise classification accuracy among randomly selected FP_11_-tagged subpopulations. For each subpopulation size, minimum accuracy between all possible pairs was determined from 10^7^ random selections, with maximum values plotted. **(G)** Comparative performance analysis of 30 candidate populations (detailed in fig. S4) using both confocal microscopy (blue) and flow cytometry (orange). Analysis methodology follows that of panel (F). Data presented as mean ± SD from three independent experiments.

To generate a stable cellular platform for evaluating the FP_11_ tag library, we engineered an expression vector encoding all three FP_1-10_ fragments (CFP_1-10_, sfCherry3Csp_1-10_, and mNG3A_1-10_) separated by self-cleaving P2A peptides, followed by an IRES and iRFP670 (Fig. 5B). Through fluorescence-activated cell sorting (FACS) based on iRFP670 expression levels, we established a HeLa cell clone (HeLa^FP1-10x3_iRFP670^) expressing high levels of three FP_1-10_ fragments.

The complete library of 96 FP_11_ tags was stably expressed in HeLa^FP1-10x3_iRFP670^ cells, with resultant populations analyzed by confocal microscopy. To quantitatively assess the fluorescence distribution patterns, we implemented spherical coordinate transformation to calculate two angular parameters: Angle_CvsG_ and Angle_RvsCG_, each ranging from 0° to 90° (Fig. 5C, left). To evaluate the discriminative power of these angular parameters, we performed pairwise population analysis using selected cell populations from the library. Cell classification was achieved using an unsupervised two-component Gaussian mixture model without prior training data (Fig. 5C, right). Classification accuracy was determined by comparing model-predicted clusters against known population identities. Superior classification accuracy correlated with greater separation of populations in the Angle_CvsG_ – Angle_RvsCG_ parameter space. This analytical framework was systematically applied to all possible pairwise combinations within the 96 FP_11_-tagged cell library (Fig. 5D).

To determine the number of distinguishable cell populations, we implemented an iterative sampling approach due to the vast number of possible patterns (approximately 2×10^20^) when selecting 20 FP_11_-tagged cells from a pool of 96 (Fig. 5E). Random subsets of predetermined sizes were drawn from the 96 FP_11_-tagged cell populations, with pairwise classification accuracies extracted from the comprehensive analysis presented in Fig. 5D. For each FP_11_-tagged subset, we identified the minimum classification accuracy, reflecting the subset’s ability to distinguish the most similar cell population pairs. This sampling process was repeated 10^7^ times, and the maximum achievable minimum accuracy was plotted as a function of subpopulation size (Fig. 5F). The analysis revealed that a subset of 20 cell populations maintained robust discrimination with a minimum pairwise classification accuracy of 90%. However, expanding to 30 populations resulted in a substantial decrease in minimum accuracy to 60%. Based on these quantitative insights, we selected an initial panel of 30 candidate FP_11_ tags (Fig. 5A, 1^st^) for subsequent refinement to establish the final optimized set of 20 tags (Fig. 5A 2^nd^) that would ensure maximal discriminative power.

We conducted comprehensive evaluation of the 30 candidate populations using both confocal microscopy and flow cytometry, with three independent experimental replicates (fig. S4). The relationship between population subset size and maximum achievable minimum pairwise classification accuracy was reassessed using these orthogonal detection methods (Fig. 5G). While fluorescence microscopy demonstrated marginally superior classification accuracy, flow cytometry maintained robust discrimination capabilities, achieving >85% accuracy across a set of top 20 FP_11_ tags. The final optimized set of 20 FP_11_ tags and their comprehensive pairwise discrimination analysis is presented in Figure S5. Here, to optimize the fluorescence signal intensity of single-color reference populations, we refined the copy number composition (Fig. 5A Final): CFP2_11_(x8) and mNG3Asp_11_(x8) were implemented at maximum repeat numbers, while sfCherry3Csp_11_ was limited to (x4) to circumvent aggregation phenomena observed at higher copy numbers (fig. S6).

### High-fidelity discrimination of twenty cell populations using optimized FP_1-10_ expression platform

The FP_1-10_ expression system was also refined by replacing the iRFP670 reporter with a puromycin resistance cassette. Through stringent puromycin selection, we established a HeLa clone cell (HeLa^FP1-10x3^) expressing elevated levels of CFP2_1-10_, mNG3Asp_1-10_, and sfCherry3Csp_1-10_. The optimized set of 20 FP_11_ tags was stably expressed in HeLa^FP1-10x3^ cells, with individual cell populations analyzed by confocal microscopy. Fluorescence was visualized with distinct spectral channels: CFP2 (blue), mNG3Asp (green), and sfCherry3Csp (red). Merged images revealed unique signatures based on both color composition and intensity distribution (Fig. 6A).

**Figure 6.**
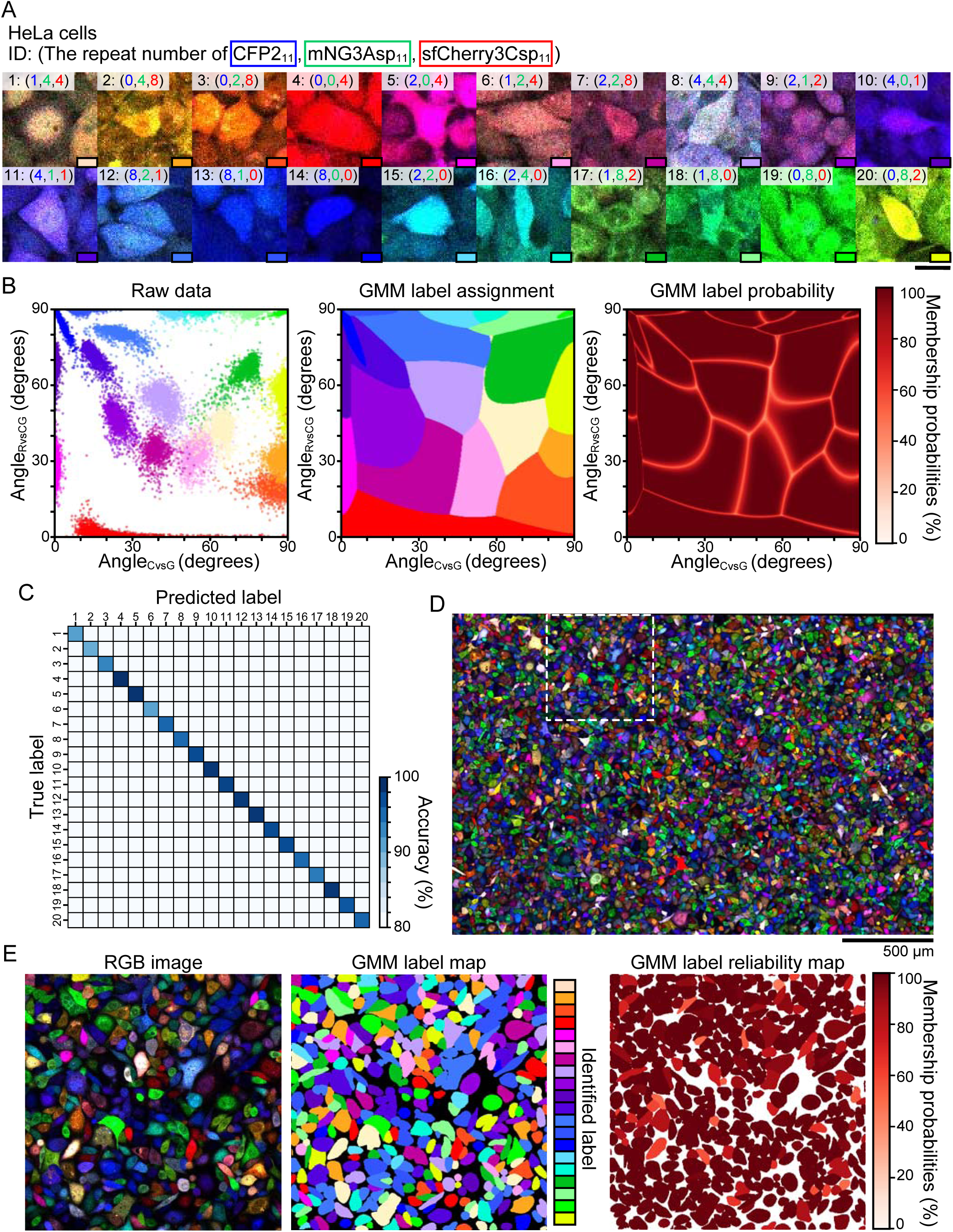
High-fidelity discrimination of twenty distinct cell populations using Caterpie. **(A)** Representative multicolor fluorescence micrographs of 20 distinct cell populations expressing unique FP_11_ tag combinations. Fluorescence channels: CFP2 (blue), mNG3Asp (green), and sfCherry3Csp (red). Copy numbers of each FP_11_ variant (CFP2_11_, mNG3Asp_11_, and sfCherry3Csp_11_) are indicated in the upper left corner of each panel. Scale bars: 30 μm. **(B)** Left: Scatter plot showing Angle_CvsG_ versus Angle_RvsCG_ distributions for 20 distinct cell populations from panel (A). (n > 1000 cells per population) Center: Distribution of label assignments in Angle_CvsG_ and Angle_RvsCG_ based on Gaussian mixture model (GMM) classification. Right: Distribution of label membership probabilities in Angle_CvsG_ and Angle_RvsCG_. **(C)** Classification performance matrix showing prediction accuracy against true population identities (rows). Color intensity indicates classification accuracy. Overall average accuracy, 97 %. **(D)** Large-field composite image of pooled cell populations from panel (A), displaying CFP2 (blue), mNG3Asp (green), and sfCherry3Csp (red) fluorescence channels. Scale bar: 500 µm. **(E)** Detailed analysis of region indicated by dashed box in panel (D). Left: Higher magnification of selected region. Center: Population assignment map following GMM-based classification into 20 distinct categories. Right: Visualization of GMM classification reliability through membership probability mapping.

Cell segmentation was performed on maximum intensity projections of composite fluorescence signals using Cellpose (*21*). Mean fluorescence intensities were quantified across all three channels for each segmented cell. Following the methodology established in Figure 5C, we implemented spherical coordinate transformation to calculate Angle_CvsG_ and Angle_RvsCG_, generating a two-dimensional angular representation of relative fluorescence intensities (Fig. 6B, left). These parameters were used to train a Gaussian mixture model (GMM), which established both label assignment distributions and label membership probability distributions in Angle_CvsG_ and Angle_RvsCG_ (Fig. 6B, center and right). Validation of the trained GMM on independent test data demonstrated exceptional classification performance, achieving 97% accuracy in discriminating all 20 cell populations (Fig. 6C). Using unsupervised k-means clustering on the 20 components achieved 96% accuracy in distinguishing between the 20 cell populations, even without requiring pre-training data for individual fluorescence profiles (fig. S7).

To evaluate system performance under practical conditions, we used pooled samples containing all 20 cell populations and acquired tiled images (Fig. 6D). Application of the trained GMM enabled robust classification of individual cells, with 78% of cells being classified with more than 95% probability (Fig. 6E).

### Interaction dynamics in multi-population cell cultures of MDCK cells expressing EGF-family ligands and their receptors

Finally, as an application of this system—named color and intensity modulation with split fluorescent proteins for cell labeling (Caterpie)—we studied the contribution of heterotypic cell interactions mediated by EGF-family ligands and their cognate receptors. For this, we used Madin-Darby Canine Kidney (MDCK) cells, which are widely used to study collective cell migration and cell competition regulated by the EGF signaling pathway. MDCK cells were labeled with the aforementioned FP_1-10_ and 20 FP_11_ tags with 96% accuracy (fig. S8A, B). Mixed cells were classified by applying a trained Gaussian mixture model (fig. S8C), 75% of cells being classified with more than 95% probability. The cell densities of twenty different labeled cell types at 72 hours post-labeling ranged within a 1.4-fold difference between the minimum and maximum values (fig. S9).

Among the four EGF-family ligands expressed in MDCK cells, we focused on HBEGF and EREG. HBEGF, a high-affinity ligand, binds to heparan sulfate proteoglycans to provide strong signals restricted to short distances, promoting localized cell migration. In contrast, EREG, a low-affinity ligand, diffuses quickly and remotely, propagating signals approximately four times faster than high-affinity ligands and more efficiently to distant cells (*22*). Both HBEGF and EREG bind to EGFR (ErbB1) and ErbB4. Using lentiviral transduction, we introduced EREG, HBEGF, EGFR (ErbB1), or ErbB4 in fluorescently labeled MDCK cells (Fig. 7A). To investigate how EGF ligands and receptors with different affinities interact between heterotypic cells and generate spontaneous patterns, these four cell types, along with wild-type cells, were co-cultured for 8 days under 3% low-serum conditions. The resulting cell population was analyzed by fluorescence microscopy to identify each cell type (Fig. 7B).

**Figure 7.**
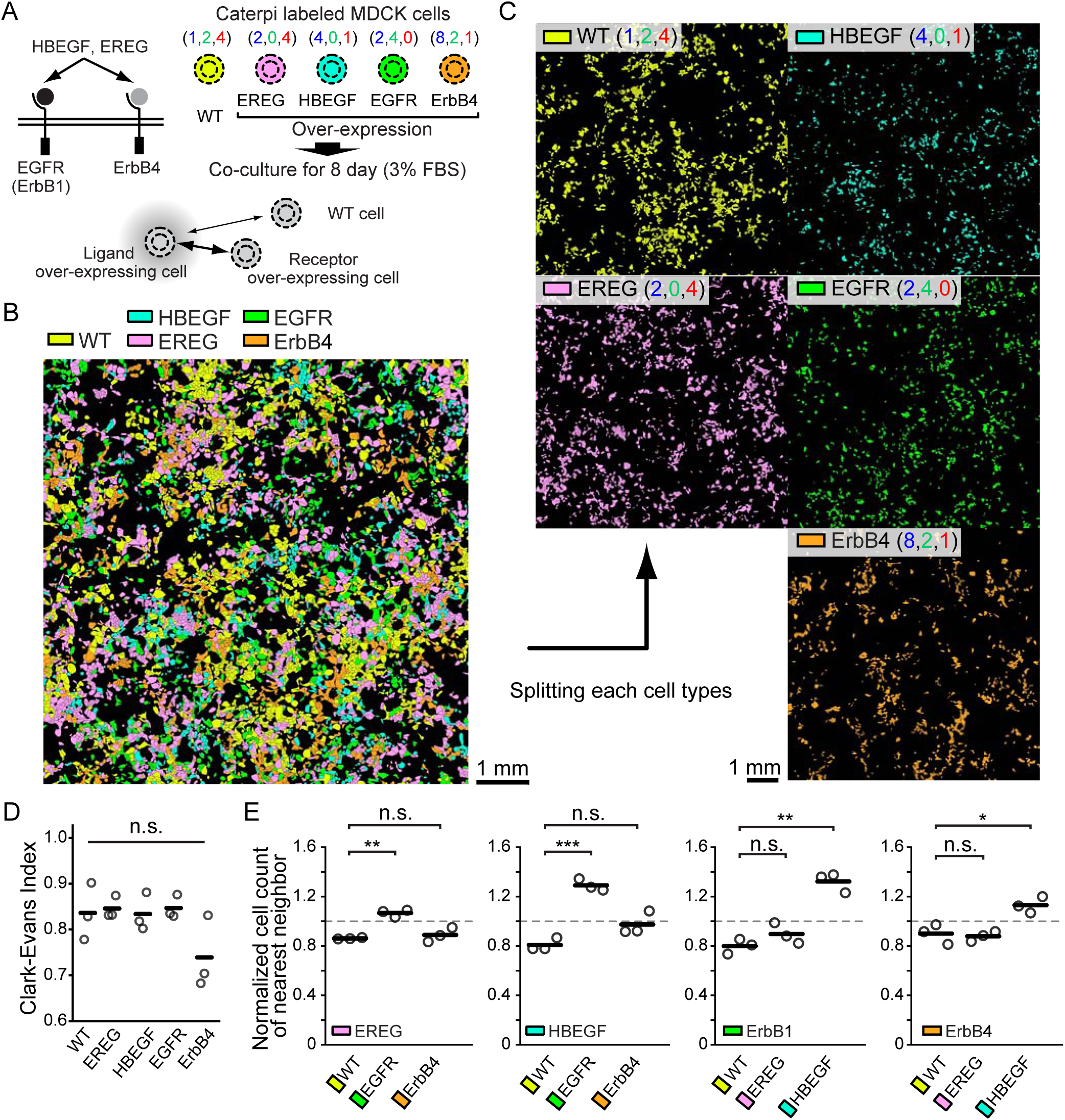
Interaction dynamics in multi-population cell cultures of MDCK cells expression EGF-family ligands and their receptors. **(A)** Schematic diagram showing the association pairs between EGF ligands (HBEGF, EREG) and EGF receptors (EGFR, ErbB2). **(B)** Representative pseudocolored image of five distinct cell types cultured together and identified using the Caterpie method. Yellow: WT, Pink: EREG, Cyan: HBEGF, Green: EGFR, Orange: ErbB4; Copy numbers of each FP11 variant (CFP211, mNG3Asp11, and sfCherry3Csp11) are (1,2,4), (2,0,4), (4,0,1), (2,4,0), and (8,2,1), respectively. Scale bar: 1 mm. **(C)** Individual display of the five cell types shown in panel E. The upper left corner of each panel indicates the re-expressed EGF signaling molecules and the copy numbers of FP_11_ tags. **(D)** Graph showing the Clark-Evans Index calculated for each of the five cell types. Data from three independent experiments. Black lines represent the mean values across the three experiments. **(E)** Normalized nearest neighbor cell analysis for evaluating spatial relationships between cells. This figure shows the ratio of heterotypic nearest neighbor counts observed in the actual data to those counted after randomly reassigning the labels of five cell types in the same dataset. Data from three independent experiments. Black lines represent the mean values across the three experiments.

Before analyzing the positional relationships between heterotypic cells, we first examined the distribution patterns of homotypic cells, as cell types with limited motility and dispersal capability affect intercellular interactions with heterotypic cells. To examine homotypic cell clustering, each cell type is individually displayed (Fig. 7C). Dispersion of each cell type was analyzed by the Clark-Evans index; values less than 1 indicate cellular clustering. We did not find significant homotypic cell clustering in any cell types (Fig. 7D). Then, we investigated the frequency of adjacent heterotypic cells. When examining receptor-expressing cells adjacent to EREG ligand or HBEGF-expressing cells (Fig. 7E), we found EGFR cells displayed significantly higher adjacency to EREG cells than wild-type cells. Conversely, when examining ligand-expressing cells adjacent to EGFR receptor or ErbB4-expressing cells, HBEGF cells exhibited higher adjacency to EGFR cells and to a lesser extent to ErbB4 in comparison to wild-type cells. These results suggest that EREG and EGF function as chemoattractants, with EREG weakly attracting distant cells, while HBEGF strongly attracts nearby cells.

In conclusion, the Caterpie method provides a versatile platform for analyzing cell populations that may exhibit homotypic and/or heterotypic clustering.

## Discussion

We have developed color and intensity modulation with split fluorescent proteins for cell labeling (Caterpie), a novel cell identification platform that achieves robust identification of 20 distinct cell populations with 96-97% accuracy. This system leverages fluorescence intensity modulation through engineered arrays of split fluorescent protein (split FP) fragments. The technological foundation of Caterpie comprises three complementary split FPs: split CFP2 and two newly engineered variants - split mNG3Asp and split sfCherry3Csp. To implement this system, we established a comprehensive toolkit consisting of 20 distinct FP_11_ tags, each constructed by fusing truncated SUMOstar protein to the 11th β-strand of the three split fluorescent proteins in varying tandem repeat configurations. These engineered FP_11_ tags demonstrate highly specific labeling of FP_1-10_-expressing cells, enabling population discrimination with 96-97% accuracy through Gaussian mixture model (GMM) analysis of the resulting multidimensional fluorescence signatures.

Current multicolor fluorescent labeling approaches can be broadly categorized into two distinct methodological frameworks. The first relies on stochastic recombination events mediated by site-specific recombinase systems such as Cre/loxP and Flp/FRT (*1*, *2*, *23*–*27*). The second approach exploits the inherent randomness of transfection efficiencies or genomic integration sites (*28*–*32*). While both established approaches depend on stochastic processes to generate fluorescent protein expression patterns, Caterpie represents a paradigm shift through its implementation of rationally designed color palettes. This deterministic approach enables precise targeting of specific cell populations and lineages. The system holds particular promise for applications requiring high-fidelity identification and longitudinal tracking of distinct cell types within complex biological contexts. However, the full realization of Caterpie’s potential necessitates further optimization of delivery methodologies. Specifically, development of refined strategies is required to achieve selective incorporation of distinct FP_11_ tag variants into target cell populations while maintaining variant segregation and preventing cross-contamination.

The theoretical prediction of an 8-fold signal enhancement for FP_11_(x8) constructs relative to their monomeric counterparts was not fully realized in experimental measurements. Our studies with split CFP2 demonstrated approximately 5-fold enhancement in median fluorescence intensity (Fig. 2C). Further investigations revealed that the choice of fusion protein significantly impacts the signal amplification efficiency of octameric constructs (fig. S2C). In research using sfGFP, when x3, x4, or x7 sfGFP_11_ was fused to β-tubulin or Lamin A/C, length and brightness correlated well, nearly achieving theoretical limits (*4*, *6*). However, when Teneurin-m was fused to x7 sfGFP_11_, only a 4-fold increase in brightness was achieved (*9*). With the goal of reducing gene size, which is an advantage of our approach, we changed from the initially used Histone H2B to ΔSUMOstar and achieved almost the same amplification rate as Histone H2B, x1.6 by each repetition (fig. S2). This collective evidence suggests that achieving linear brightness amplification through multimerization of the 11^th^β-strand requires careful optimization of the fusion partner.

The application of computational protein structure prediction proved instrumental in enhancing split fluorescent protein performance (Fig. 3). This approach represents a significant departure from conventional optimization strategies, which typically rely on random mutagenesis (*33*). Simultaneous introduction of mutations in both β_1-10_ and β_11_ regions that enhance their mutual interactions proved challenging through random mutagenesis, because the statistical probability of obtaining cooperative mutations is exceedingly low. Our successful implementation of structure prediction-guided optimization suggests a broader applicability of this methodology across other split fluorescent protein systems.

The discriminative capacity of the Caterpie system holds significant potential for expansion through two primary avenues: the development of novel split fluorescent protein variants and the integration of subcellular targeting sequences. Furthermore, compatibility with the Landing Pad system (*34*), which enables precise single-copy genomic integration, presents opportunities for multiplexed cellular labeling. These technological developments collectively position Caterpie as a versatile platform for investigating complex biological processes requiring simultaneous tracking of multiple distinct cell populations.

## Materials and Methods

### Plasmids

The following cDNAs were synthesized with optimized codons by GeneArt (Thermo Fisher Scientific, Waltham, MA): sfGFP_1-10_ (*3*), CFP2_1-10_ (*35*), mNeonGreen3A_1-10_ (*11*), sfCherry3Csp_1-10_, mRuby4_1-10_ (*10*), GB1 (*18*), ΔSUMOstar (*19*). Additional split fluorescent protein variants were generated through site-directed mutagenesis of existing templates: Cerulean_1-10_ (*10*) and YFP1_1-10_ (*35*) from sfGFP_1-10_, mNeonGreen3K_1-10_ (*11*) from mNeonGreen3A_1-10_, and sfCherry3C_1-10_ (*12*) from sfCherry2_1-10_ (Addgene: #82603). The Histone H2B coding sequence was subcloned from Addgene (plasmid # 11680 for Histone H2B; Cambridge, MA).

### Vector Construction for Stable Cell Line Generation

A base Tol2 transposon vector (pT2A-IRES-iRFP670) was constructed by subcloning iRFP670 cDNA (*36*) with an internal ribosome entry site (IRES) (*37*) into the pT2A vector (*38*). CFP2_1-10_ cDNA was subsequently PCR-amplified and inserted into this base vector using In-Fusion assembly (Clontech, Mountain View, CA) to generate pT2A_CFP2_1-10_-IRES-iRFP670. Two distinct vector backbones, pT2A_IRES-iRFP670 and pT2ADW (containing IRES-puro cassette) (*39*), were used for multicistronic construct assembly. The following elements were PCR-amplified and assembled into both vectors using In-Fusion: sfCherry3Csp_1-10_, mNeonGreen3Asp_1-10_ with self-cleaving P2A peptide (*40*), and CFP2_1-10_ with P2A peptide. Stable cell lines were generated through co-transfection of the constructed pT2A vectors with pCS-TP (obtained from Kawakami et al., 2004 (*38*)).

To generate pPBpuro_EF1a constructs containing CFP2_11_(x1), CFP2_11_(x2), CFP2_11_(x4), CFP2_11_(x8) -mCherry-NLS, cDNAs encoding Histone H2B, self-cleaving peptide P2A sequences, mCherry, and the nuclear localization signal (NLS) of the SV40 large T antigen (PKKKRKV) (*41*) were PCR-amplified and assembled using In-Fusion into pPBpuro_EF1a vectors (a kind gift from K. Yusa), yielding pPBpuro_EF1a_Histone H2B-P2A-mCherry-NLS. Either synthesized CFP2_11_ or CFP2_Full-length_ cDNA (created by PCR amplification of CFP2_1-10_ cDNA and CFP2_11_) was then assembled into pPBpuro_EF1a_Histone H2B-P2A-mCherry-NLS, resulting in pPBpuro_EF1a_CFP2_Full-length_/CFP2_11_(x1)-mCherry-NLS. Restriction enzyme sites (BglII before FP_11_, BamHI and NotI after FP_11_) were introduced to facilitate the construction of tandem repeats. To generate CFP2_11_(x2), the CFP2_11_(x1) insert and vector were digested with BglII/NotI and BamHI/NotI, respectively, and then ligated. Subsequently, CFP2_11_(x4) and CFP2_11_(x8) were generated using the same approach. By substituting the CFP2_11_ insert with sfCherry3Csp_11_ or mNeonGreen3Asp_11_ inserts, multiple tandem repeats of these fragments were constructed using the same strategy. Various modifications were made to customize the constructs for specific experimental requirements: the puromycin-resistance gene (puro) was replaced with the blasticidin S-resistance gene (bsr); Histone H2B was substituted with GB1, ΔSUMOstar; mCherry was replaced with EBFP2 or mTagBFP2; and the NLS was substituted with the nuclear export signal (NES) of the HIV-1 rev protein (LPPLERLTLD) (*42*).

Mutant variants of sfCherry3C_1-10_ were generated using overlap extension PCR and assembled into pPB-based vectors (*43*) containing IRES-bsr (blasticidin S-resistance gene) using In-Fusion. Additional mutant variants of mNeonGreen3A_11_ and sfCherry3C_11_ were synthesized and inserted into pPB-based vectors (*43*) containing IRES-puro (puromycin-resistance gene) using Ligation High Ver. 2 (TOYOBO). Stable cell lines were established through co-transfection of these constructs with pCMV-mPBase (obtained from the Wellcome Trust Sanger Institute).

### Cell culture and establishment of cell lines

HeLa cells and Lenti-X 293T cells were obtained from the Human Science Research Resources Bank and Clontech, respectively. Both cell lines were maintained in DMEM (Wako Pure Chemical Industries) supplemented with 10% fetal bovine serum (Sigma-Aldrich) and 1% penicillin/streptomycin (Nacalai Tesque). MDCK (ECACC 84121903) cells were purchased from the European Collection of Authenticated Cell Cultures (ECACC) through the RIKEN BioResource Center (no. RCB0995) and maintained in DMEM (Wako Pure Chemical Industries) supplemented with 10% fetal bovine serum (Sigma-Aldrich), 1% penicillin/streptomycin (Nacalai Tesque).

For transposon-mediated gene transfer, pT2A_CFP2_1-10_-IRES-iRFP670, or pT2A_FP_1-10_x3-IRES-iRFP670 was cotransfected with pCS-TP into HeLa cells by using 293fectin (Thermo Fisher Scientific, Waltham, MA). These obtained HeLa cells were sorted using an FACS Aria IIu cell sorter (Becton Dickinson, Franklin Lakes, NJ) based on iRFP670 fluorescence to achieve a high expression level of the CFP2_1-10_ or FP_1-10_x3. Single-cell cloning of these sorted populations to yield HeLa^CFP2^ ^1-10_iRFP670^ cells and HeLa^FP1-10x3_iRFP670^ cells.

pT2A_FP_1-10_x3-IRES-iRFP670 was cotransfected with pCS-TP using either 293fectin (Thermo Fisher Scientific, Waltham, MA) for HeLa cells or electroporation with an Amaxa nucleofector (Lonza, Basel, Switzerland) for MDCK cells. The transfected cells were selected with 5 μg mL^-1^ puromycin (no. P-8833; Sigma-Aldrich), followed by single-cell cloning to yield HeLa^FP1-10x3^ cells or MDCK^FP1-10x3^ cell lines.

### Fluorescence imaging

For evaluation of split fluorescent proteins, HeLa cells in 24-well plates were co-transfected with 100 ng of FP_11_ plasmid [FP_Full-length_, FP_11_(x1), or FP_11_(x4)] and 400 ng of FP_1-10_ plasmid using 293fectin (Thermo Fisher Scientific, Waltham, MA). Cells were seeded onto collagen type I-coated (Nitta Gelatin, Osaka, Japan) glass-based 24-well plates (AGC Inc., Tokyo, Japan) and cultured for 48 hours. Prior to imaging, cells were equilibrated for at least 1 h in DMEM/F12 (Thermo Fisher Scientific) supplemented with 10% fetal bovine serum (Sigma-Aldrich), and penicillin/streptomycin (Nacalai Tesque).

For FP_11_ labeling,

pPBbsr_EF1a-ΔSUMOstar-CFP2_11_(xa)-mNeonGreen3Asp_11_(xb)-sfCherry3Csp_11_(xc) was cotransfected with pCMV-mPBase into HeLa^FP1-10x3^ cells by using 293fectin or MDCK^FP1-10x3^ cells using electroporation. Transfected cells were selected with 10 μg mL^-1^ blasticidin S (Calbiochem, San Diego, CA) and seeded onto collagen type I-coated glass-based 24-well plates.

Cells were observed with a Leica TCS SP8 FALCON confocal microscope (Leica-Microsystems, Wetzlar, Germany) equipped with an HC PL APO 20x/0.75 dry CS2 objective, an HC PL APO 40x/1.30 OIL CS2 objective, Leica HyD SMD detectors, a white light laser of 80 MHz pulse frequency, a Diode 405 (VLK 0550 T01; LASOS, Jena, Germany), a 440 nm diode laser (PDL 800-D; PicoQuant, Berlin, Germany), and a stage top incubator (Tokai Hit, Fujinomiya, Japan) to maintain 37 °C and 5% CO2. The following excitation wavelengths and emission band paths were used for the imaging: for EBFP2 imaging, 405 nm excitation, 410–450 nm emission; Cerulean or CFP2 imaging, 440 nm excitation, 460–490 nm emission; for sfGFP imaging, 488 nm excitation, 500–550 nm emission; for YFP1 imaging, 514 nm excitation, 520–550 nm emission; for mNG3A, mNG3K, or mNG3Asp imaging, 506 nm excitation, 510–550 nm emission; for sfCherry3C or sfCherry3Csp imaging, 594 nm excitation, 600–645 nm emission; for mRuby4 imaging, 561 nm excitation, 580–645 nm emission; for iRFP670 imaging, 650 nm excitation, 660–760 nm emission. To eliminate the background signal, the time gate for fluorescence detection was set from 1.0 ns to 6.0 ns.

In the multiplex imaging, the following excitation wavelengths and emission band paths were used: for CFP2 imaging, 440 nm excitation, 450–500 nm emission; for mNG3Asp imaging, 506 nm excitation, 515–580 nm emission; for sfCherry3Csp imaging, 594 nm excitation, 605–645 nm emission. To eliminate the background signal, the time gate for mNG3Asp and sfCherry3Csp fluorescence detection was set from 0.2 ns to 6.0 ns.

Images were processed and analyzed with ImageJ (*44*). For quantification of fluorescent intensity, images were segmented using Cellpose algorithm (*21*). Fluorescent intensity calculation was performed using custom MATLAB (MathWorks) and Python scripts.

### Fluorescence spectral analysis

Fluorescence spectra were measured using a Leica TCS SP8 FALCON inverted microscope. Images were acquired with an HC PL APO 20x/0.75 dry CS2 objective using excitation at either 470 nm for split mNG3A or 546 nm for split sfCherry3C.

### Flow cytometry analysis

Cells were suspended in PBS containing 3% FBS and analyzed or sorted with a FACS Aria IIu cell sorter (Becton Dickinson, Franklin Lakes, NJ). The following laser and emission filter combinations were used for fluorescence detection: CFP2, a 407 nm laser and an ET470/24m filter (Chroma Technology Corp., Bellows Falls, VT); mNeonGreen3Asp, a 488 nm laser, and a DF530/30 filter; sfCherry3Csp, a 561 nm laser and a DF582/15 filter (Omega Optical); iRFP670, a 633 nm laser and a DF660/20 filter (Omega Optical). Cell debris and aggregates were excluded by gating for size and granularity. Data analysis was performed using FlowJo software (Tree Star, Ashland, OR).

### Lentivirus infection

For lentivirus production, the EGF ligand or receptor expressing plasmid, psPAX2 (Addgene no. 12260), and pCMV-VSV-G-RSV-Rev (*45*) were co-transfected into Lenti-X 293T cells using polyethylenimine (no. 24765-1; Polyscience Inc.). The infected cells were selected with media containing the following antibiotics, depending on the drug resistance genes carried by the EGF ligand or receptor expressing plasmids: 200 μg ml^−1^ hygromycin (no. 31282-04-9; Wako).

### Cell growth

For quantifying cell growth, MDCK cells labeled by Caterpie were seeded on collagen-coated 24-well glass-bottom plates (AGC Inc., Tokyo, Japan) at a density of 5×10^4^ cells/ml. After 1 hour of incubation, the medium was replaced with DMEM/F12 supplemented with 100 units/ml penicillin, 100 μg/ml streptomycin, and 10% FBS. Following cell seeding, cells were observed using a confocal microscope (Leica SP8) every 24 hours for 3 days. For observation conditions, please refer to the "Fluorescent imaging" section. Cell numbers were counted using the Cellpose segmentation algorithm.

### Computational design for enhanced β-barrel stability and split-strand complementation

The tertiary structures of split mNG3Asp and split sfCherry3Csp were modeled using ColabFold (13) implementation of AlphaFold Multimer (14) or AlphaFold2 (17) and visualized with PyMOL (http://www.pymol.org/). The highest-confidence model of the split sfCherry3Csp was then adopted as the starting coordinate for downstream Rosetta-based design.

To enhance global β-barrel stability of split sfCherry3Csp, *in silico* site-saturation mutagenesis (SSM) of all 224 positions in sfCherry3C□–□□ and sfCherry3C□□ was performed with a Rosetta-based workflow adapted from Thieker et al (16). For each variant, side-chain and backbone sampling were confined to a 10□Å sphere around the mutated residue, while the remainder of the protein was held fixed under coordinate constraints (sd = 1 outside the design sphere; sd = 2 within a surrounding “soft sphere”). Each mutant underwent three cycles of Cartesian FastRelax (MonomerRelax2019), and the change in Rosetta energy (ΔE) was calculated as the difference in total_score between each mutant and the wild-type model using the ref2015 score function.

To reinforce β□–□□/β□□ interface, two design workflows, Point-Mutation (PM) and Mutation Cluster (MC), were adapted from the workflow of Thieker et al. and modified for interface optimization. (1) In the PM workflow, SSM of the β□□ fragment (residues 1–16) was carried out, with the addition of an InterfaceAnalyzer to compute binding energy (dG_separated) and packing statistics. Residues within 10 Å of each mutation site were sampled under the same constraint scheme used for stability design, while non-neighbor residues were held fixed; each point mutant was relaxed by Cartesian FastRelax prior to interface analysis, and final dG_separated values were calculated using the unconstrained ref2015 Rosetta score function. (2) In the MC workflow, the seed mutations from the PM step were proceeded into a FastDesign protocol using the InterfaceDesign2019 relax script to generate clustered mutations. The neighbor-selection radius was extended to 12 Å, and an inner design shell was defined by residues both in direct atomic contact (≤ 5–7 Å, via a CloseContact selector) and satisfying the InterfaceByVector geometric filter. A ResidueTypeConstraintGenerator was applied to inner-shell, biasing retention of wild-type identities. Critical chromophore-interacting residues were protected by marking them non-designable. Multi-residue combinations sampled via FastDesign, and each cluster variant was evaluated with InterfaceAnalyzer for dG_separated and packing quality. The scores were obtained under the unconstrained ref2015 function.

### Calculation of angular data from individual cells

Three-dimensional fluorescence intensity data were acquired from individual cells by measuring CFP2, mNG3Asp, and sfCherry3Csp signals with background subtraction. These measurements were represented in a 3D coordinate system, with CFP2 intensities plotted on the x-axis, mNG3Asp on the y-axis, and sfCherry3Csp on the z-axis. For each cell, the Cartesian coordinates (x, y, z) were transformed into spherical coordinates (r, Angle_CvsG,_ Angle_RvsCG_) using a custom-made Python script. The coordinate transformation was performed using standard spherical coordinate conversion equations.

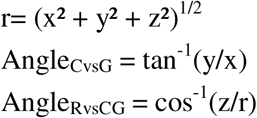

### Two-component Gaussian mixture model (GMM) for FP_11_ tag screening

Fluorescence intensity measurements from individual cells across 96 distinct cell populations were transformed into angular coordinates (Angle_CvsG_ and Angle_RvsCG_). To enhance clustering robustness, outliers were removed by excluding the top and bottom 10% of data points that exhibited the greatest deviation from their respective cluster centers in both angular measurements. An unsupervised two-component GMM was implemented using scikit-learn (version 0.24.2; Pedregosa et al., 2011 (*46*)) to perform pairwise population comparisons. For random sampling, a predetermined number of cell populations were selected from a total of 96 populations using simple random sampling without replacement. All computational analyses were performed using custom Python scripts.

### Cell classification using Gaussian mixture model (GMM)

Prior to angular calculations, we subtracted background and normalized the fluorescence intensities of CFP2, mNG3Asp, and sfCherry3Csp by dividing each value by the median intensity across all 20 cell types. Using these normalized fluorescence values, we then calculated Angle_CvsG_ and Angle_RvsCG_ measurements for subsequent analysis.

The classification model was trained using GMMs implemented in the scikit-learn library. we first fitted a single-Gaussian distribution to the training subset of each class individually; the resulting class-specific means and covariance matrices were then concatenated to obtain the GMM. The training dataset comprised 80% of the total data (∼2.9x10^4 cells in total, n > 10^3 cells per cell population), consisting of two-dimensional features (Angle_CvsG_ and Angle_RvsCG_ measurements) from 20 distinct cell populations. The trained GMMs were then applied to angular data from pooled cells, and the predicted cell labels were visualized. For each class k, we calculated the probability of each input data point belonging to the corresponding GMM model, representing the likelihood of class membership:

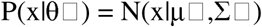

where x is the input data point, θ□ represents the parameters of the k-th GMM model, and N(x|μ□,Σ□) denotes the normal distribution with mean μ□ and covariance Σ□. Subsequently, the probabilities were normalized such that the sum of probabilities across all classes equals 1 for each data point:

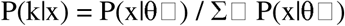

We visualized the membership probabilities of the predicted classes. All computational analyses were performed using custom-made Python scripts.

### Cell classification using K-means

We used Angle_CvsG_ and Angle_RvsCG_ as the feature space coordinates for clustering. K-means clustering was performed on the resulting dataset using scikit-learn’s KMeans algorithm with 21 clusters (random_state = 0). As shown in Figure 6B, cells labeled with tag (BGR)=(0,0,4) corresponding to ID #4 exhibited a wide distribution of values in the Angle_CvsG_-Angle_RvsCG_ space, resulting in their identification as two distinct clusters that were subsequently merged. This accounts for the selection of 21 clusters in our analysis. The predicted cluster labels were then remapped to match the original cell population labels for validation. Classification accuracy was assessed by comparing the remapped cluster assignments with ground truth labels and visualized using a confusion matrix. The overall classification accuracy was calculated as the percentage of correctly classified cells. All calculations were performed using Python.

### Clark-Evans Index

We analyzed the spatial distribution patterns of five cell types. Individual cells were first segmented using Cellpose, with centroids calculated and classified into five cell types via the Caterpie method. For spatial analysis, we extracted the coordinates of each cell type, calculated nearest neighbor distances using a k-dimensional tree algorithm, and computed the Clark-Evans index (R). This index compares observed mean distances to those expected under complete spatial randomness, where R = 1 indicates random distribution, R < 1 indicates clustering, and R > 1 indicates regularity. All calculations were performed using Python.

### Cell count of the nearest neighbor

We analyzed the spatial relationships between different cell types using a nearest-neighbor approach. Cells were classified into five distinct populations based on fluorescent labels. For each cell of the target population (label 1), we calculated the distance to every cell of other populations (labels 2-5) and identified the closest neighboring cell using a k-dimensional tree algorithm. To assess the statistical significance of the observed patterns, we generated 100 randomized distributions for each dataset by shuffling cell positions while preserving the total number of each cell type. To determine whether the observed spatial associations between cell types differed significantly from random expectations, we calculated the ratio of the frequency with which each non-target cell type appeared as the nearest neighbor to target cells in the original distribution divided by the mean frequency observed across 100 randomized distributions. All calculations were performed using Python.

### Statistics

All statistical analyses were carried out using Python. No statistical analysis was used to predetermine the sample size. An unpaired Welch’s t-test was used to evaluate statistically significant differences. Data are expressed as the mean□±□s.d. *P*-values of less than 0.05 were considered to be statistically significant in two-tailed tests and were classified into four categories: \**p* < 0.05, \*\**p* < 0.01, \*\*\**p* < 0.001, and n.s. (not significant, i.e., *p* ≥ 0.05).

## Figure Legend

**Supplementary Figure S1.**
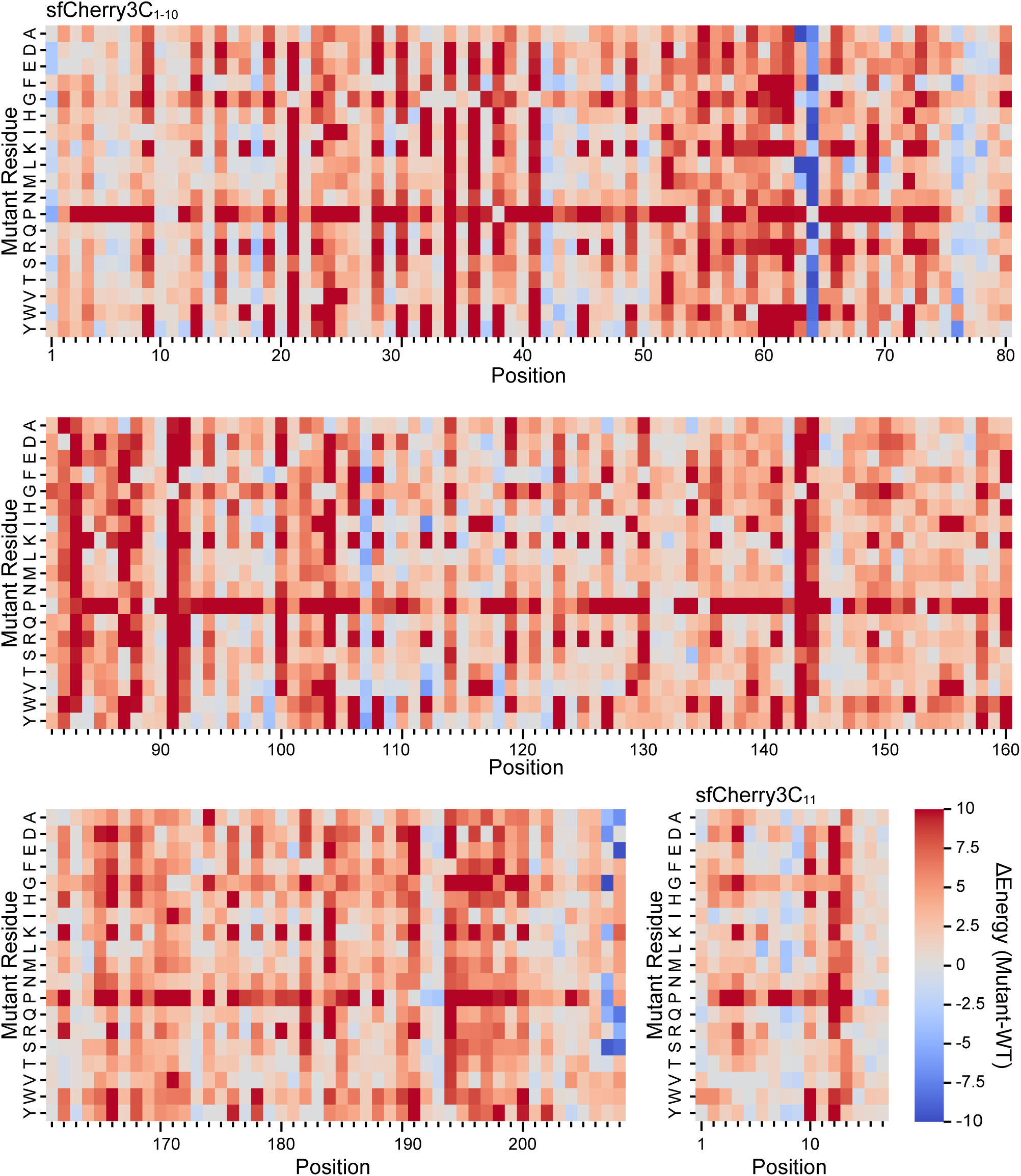
Comprehensive mutational scanning of split sfCherry3C. Heatmap displaying the ΔEnergy values for saturating mutations in both sfCherry_1-10_ and sfCherry_11_.

**Supplementary Figure S2.**
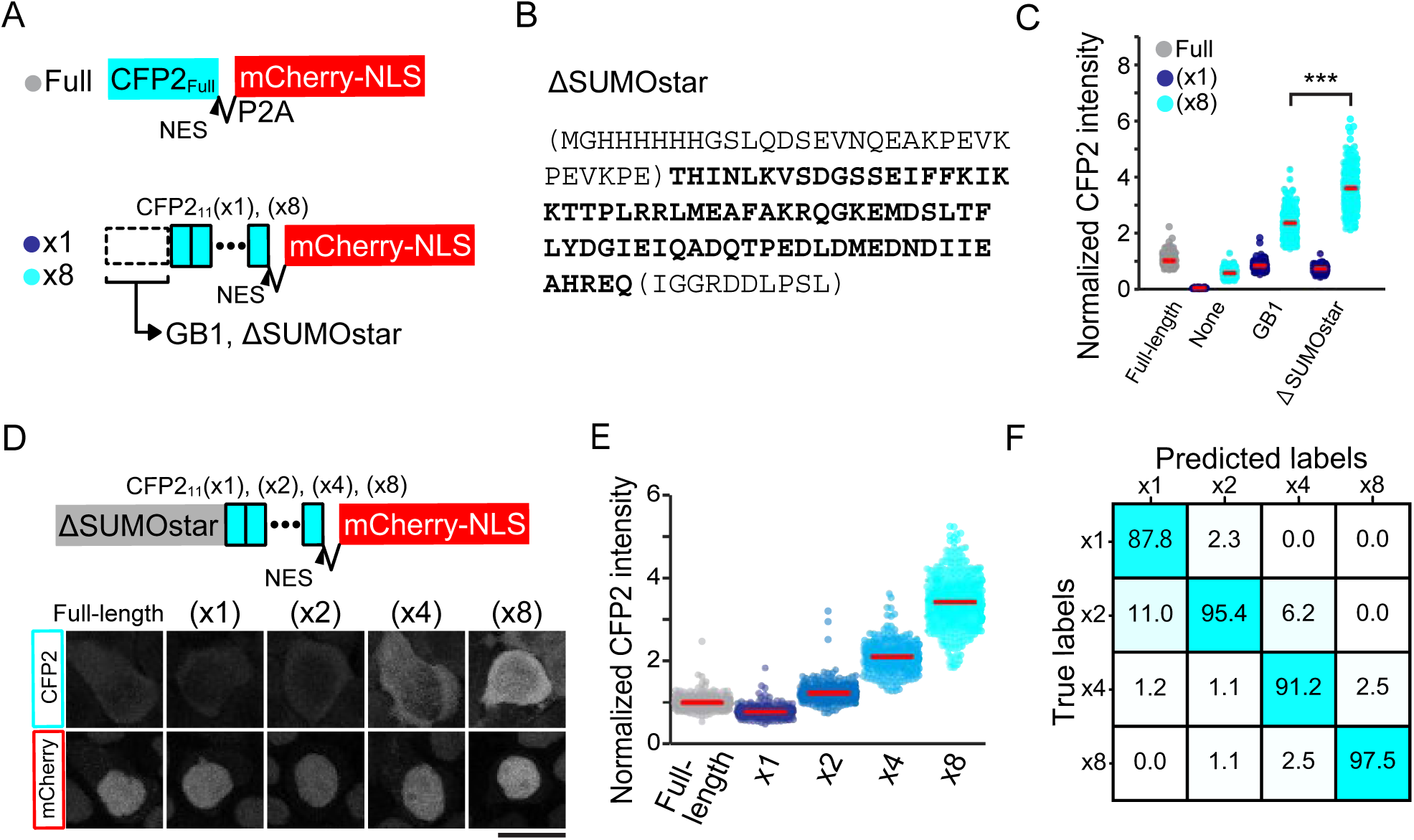
Optimization of signal amplification using truncated SUMOstar fusion strategy. **(A)** Schematic representation of expression constructs: CFP2_Full-length_-NES and CFP2_11_ variants [(x1) or (x8)]-NES with or without GB1 or ΔSUMOstar (truncated SUMOstar) tags. **(B)** Amino acid sequences of SUMOstar, with ΔSUMOstar region highlighted in bold. **(C)** Bee swarm plots showing quantitative analysis of normalized CFP2 fluorescence in HeLa cells co-expressing CFP2_1-10_ with either CFP2_Full-length_-NES or CFP2_11_(x1)/CFP2_11_(x8)-NES variants, with or without GB1 or ΔSUMOstar tags (constructs shown in panel A). Median values are indicated by red lines. **(D)** Construct architecture and expression analysis: Upper panel shows schematics of ΔSUMOstar fusions containing varying copy numbers of CFP2_11_ [CFP2_11_(x1), (x2), (x4), and (x8)]. Lower panel presents representative confocal micrographs of HeLa cells stably co-expressing CFP2_1-10_ with either CFP2_Full-length_-NES or ΔSUMOstar-tagged CFP2_11_ variant. Scale bar: 20□µm. **(E)** Comparative analysis of normalized CFP2 fluorescence in HeLa cells expressing ΔSUMOstar-tagged CFP2_11_ variants [(x1), (x2), (x4), or (x8)] shown in panel D. Data presented as bee swarm plots with median values indicated by red lines. **(F)** Classification performance matrix showing predictive accuracy against true population identities (rows), with color intensity indicating classification accuracy.

**Supplementary Figure S3.**
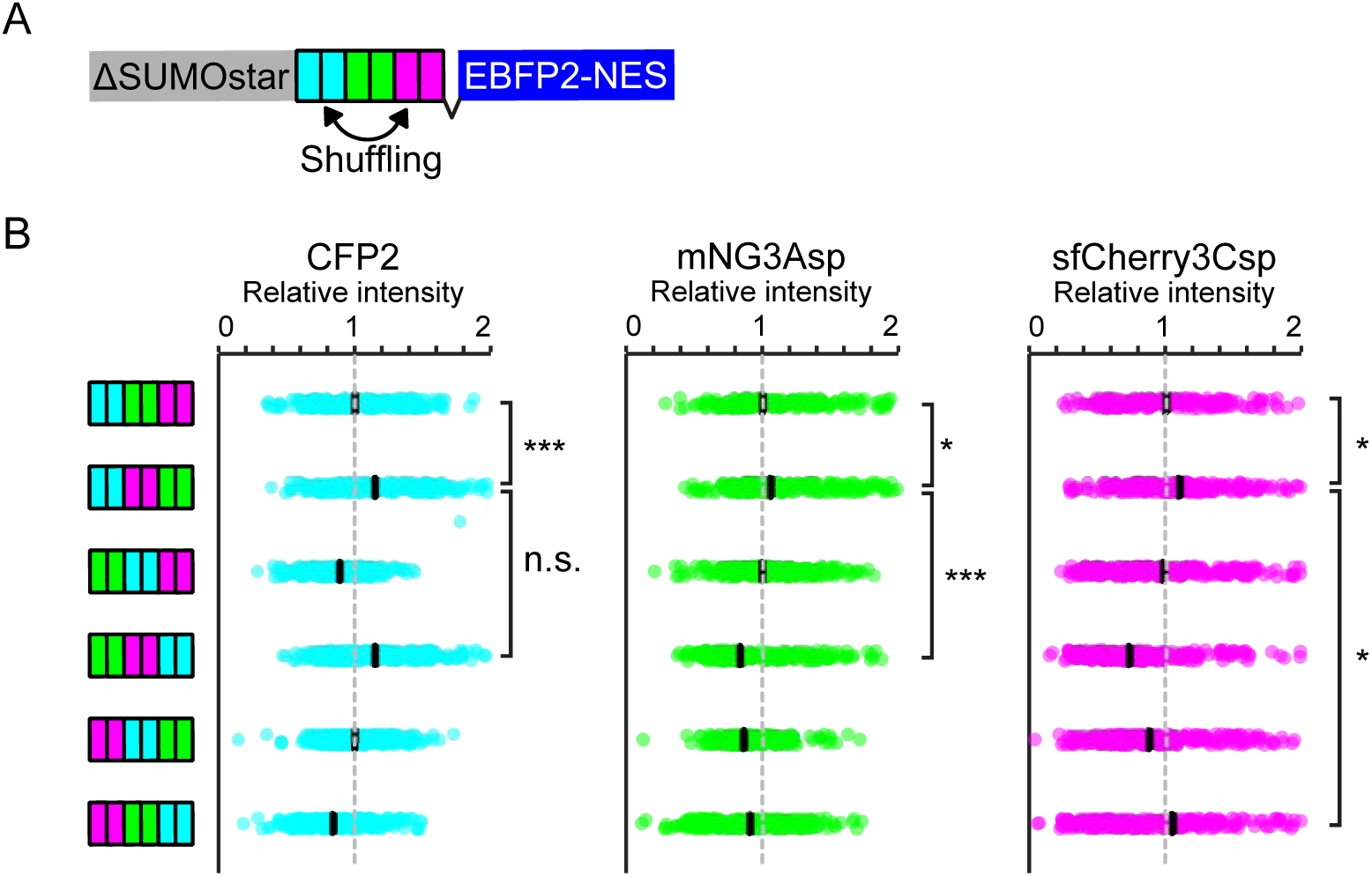
Influence of FP_11_ fragment order on fluorescence signal distribution. **(A)** Schematic representation of expression constructs: ΔSUMO-tagged FP_11_ arrays comprising paired repeats of CFP2_11_, mNG3Asp_11_, and sfCherry3Csp_11_ [(x2) each, total x6]. **(B)** Quantitative analysis presented as bee swarm plots showing normalized fluorescence intensities (CFP2, mNG3Asp, and sfCherry3Csp) in HeLa cells expressing six different permutations of FP_11_ repeat arrangements, with concurrent FP_1-10_x3 expression. Data obtained by confocal microscopy, with median values indicated by black lines.

**Supplementary Figure S4.**
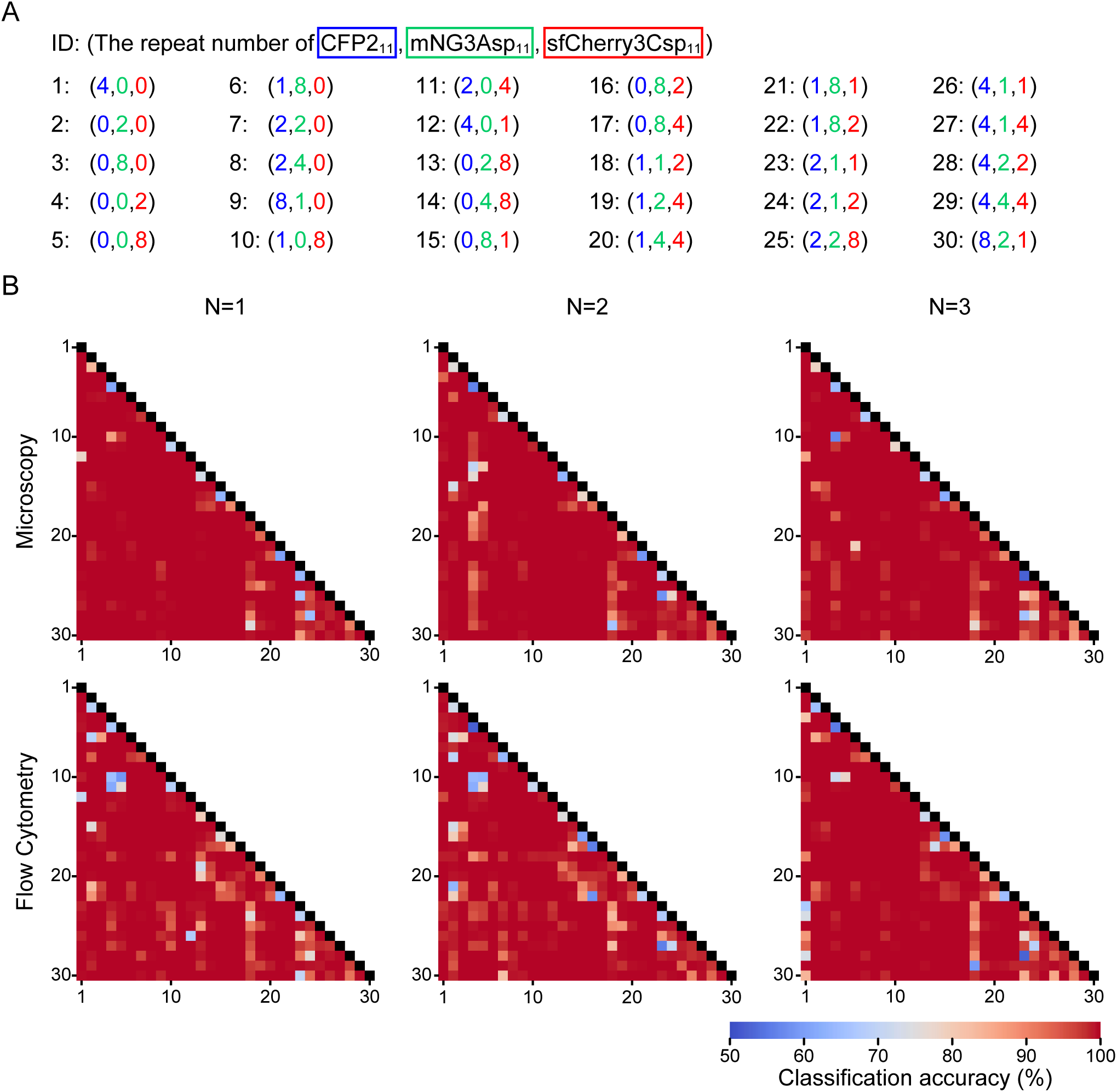
Comparative analysis of binary classification performance across 30 selected cell populations. **(A)** Catalog of 30 selected FP_11_ tag variants (detailed in Fig. 5A), detailing identification numbers and copy numbers of CFP2_11_, mNG3Asp_11_, and sfCherry3Csp_11_ fragments. **(B)** Matrix analysis of pairwise classification accuracies for 30 cell populations expressing distinct FP_11_ tag combinations (listed in panel A), evaluated by both microscopy (upper panel) and flow cytometry (lower panel). Data represent results from three independent experiments.

**Supplementary Figure S5.**
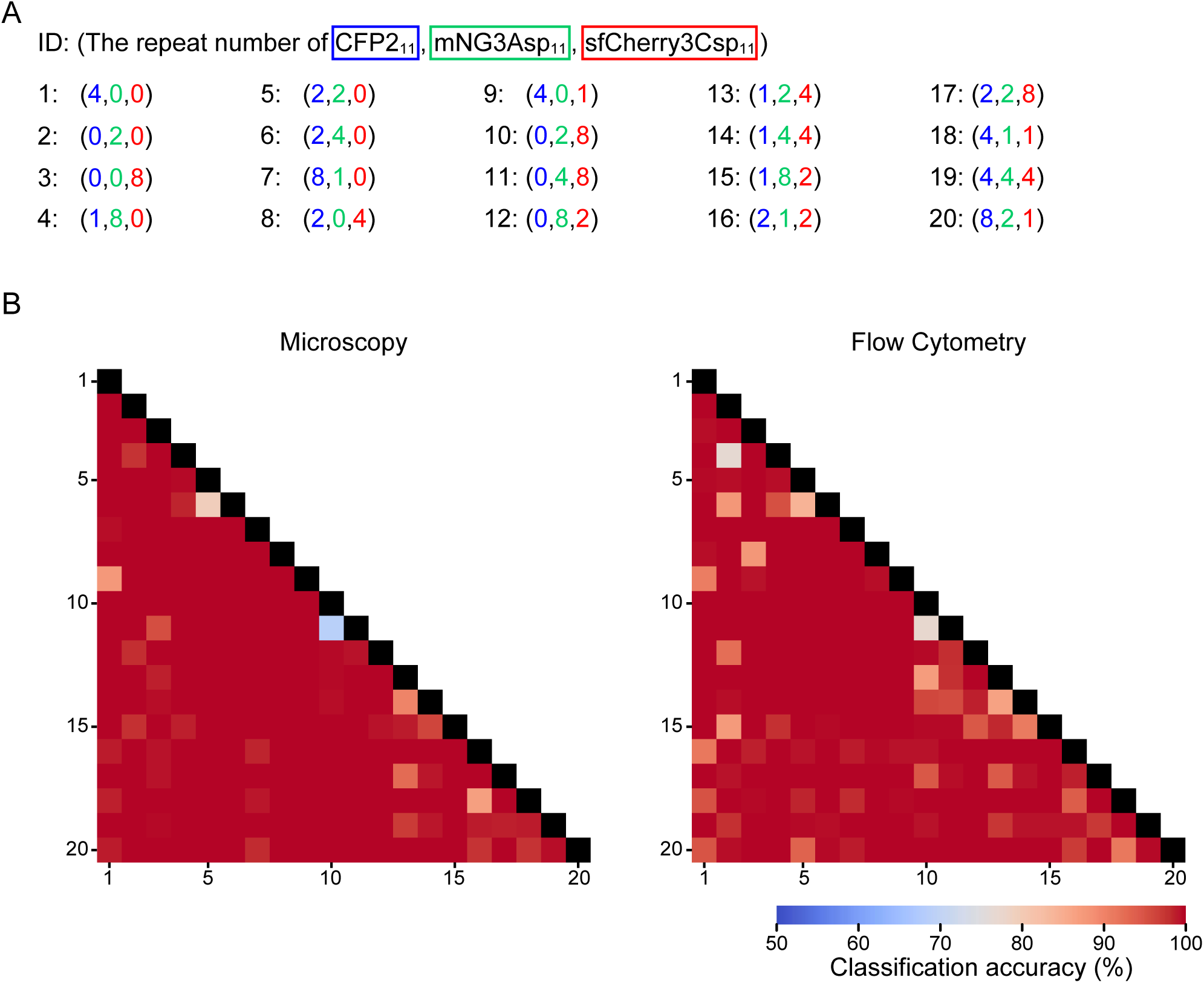
Binary classification performance analysis of final 20 cell population panel. **(A)** Catalog of optimized FP_11_ tag set comprising 20 selected variants (detailed in Fig. 5A), detailing identification numbers and copy numbers of CFP2_11_, mNG3Asp_11_, and sfCherry3Csp_11_ fragments. **(B)** Matrix analysis of pairwise classification accuracy for 20 cell populations expressing distinct FP_11_ tag combinations (defined in panel A). Comparative evaluation performed using both microscopy (left panel) and flow cytometry (right panel). Data represent mean values from three independent experiments in Supplementary Figure S3.

**Supplementary Figure S6.**
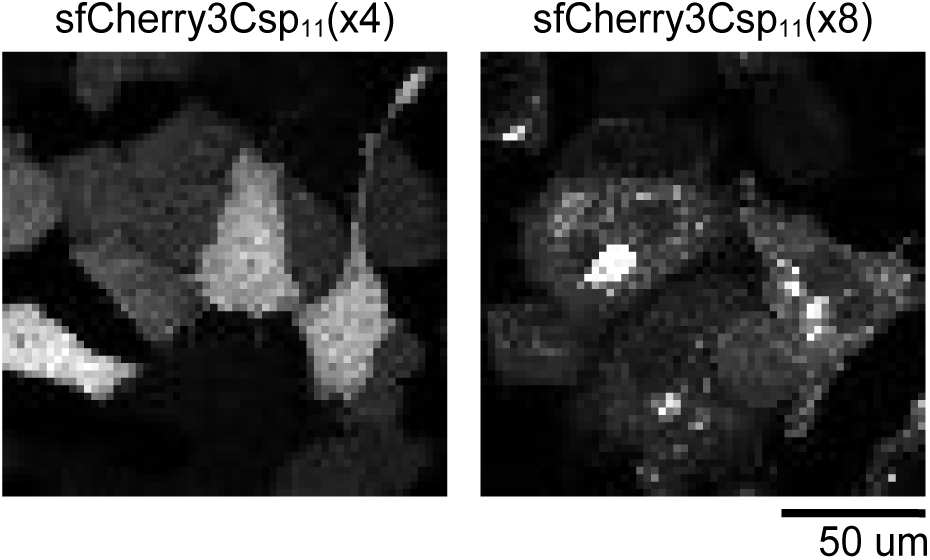
Aggregation character of sfCherry3Csp_11_(x8) Representative images of sfCherry3Csp fluorescence in HeLa cells expressing ΔSUMOstar-tagged sfCherry3Csp_11_ variants [(x4), or (x8)], with concurrent FP_1-10_x3 expression.

**Supplementary Figure S7.**
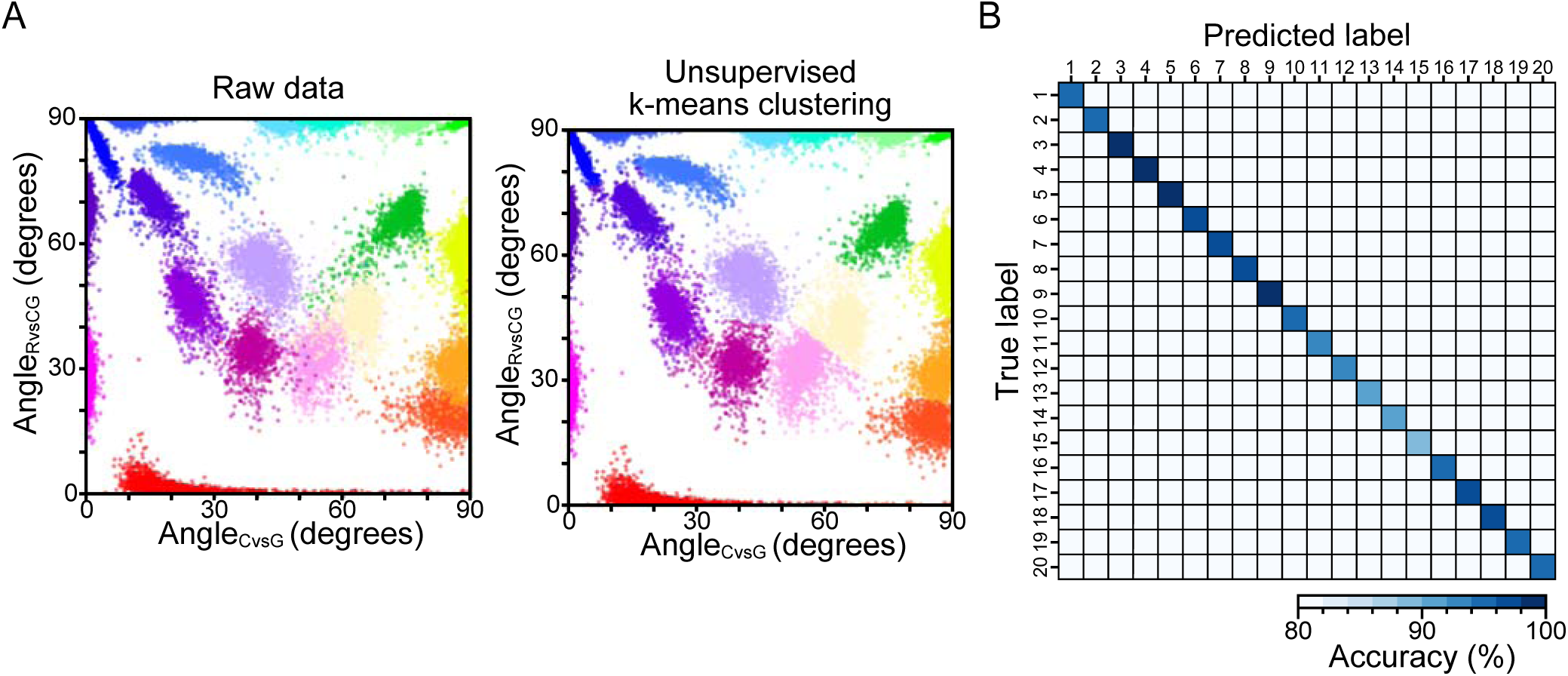
Unsupervised k-means clustering of 20 cell populations. **(A)** Left: Scatter plot showing the distribution of Angle_CvsG_ versus Angle_RvsCG_ for 20 distinct cell populations (n > 1,000 cells per population). Right: Scatter plot showing the distribution of Angle_CvsG_ versus Angle_RvsCG_ for individual cells grouped into 20 distinct populations using unsupervised k-means clustering. **(B)** Classification performance matrix of unsupervised k-means classification in panel (A). Color intensity indicates classification accuracy. Overall average accuracy, 96 %.

**Supplementary Figure S8.**
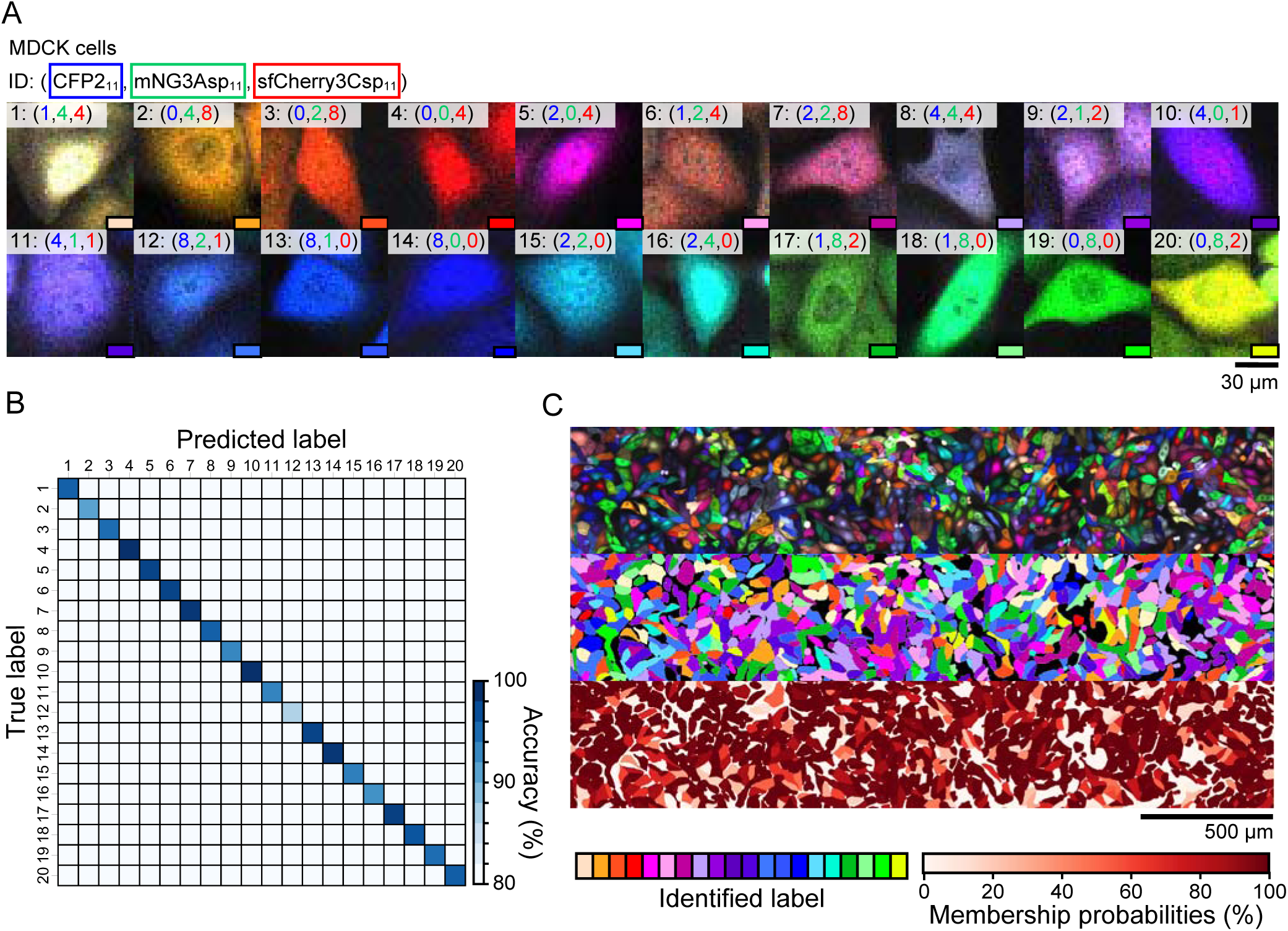
High-fidelity discrimination of twenty distinct cell populations in MDCK cells using Caterpie. **(A)** Representative multicolor fluorescence micrographs of 20 distinct cell populations expressing FP_11_ tags. Fluorescence channels: CFP2 (blue), mNG3Asp (green), and sfCherry3Csp (red). Copy numbers of each FP_11_ variant (CFP2_11_, mNG3Asp_11_, and sfCherry3Csp_11_) are indicated in the top of each panel. Scale bars: 30 μm. **(B)** Classification performance matrix showing prediction accuracy against true population identities (rows). Color intensity indicates classification accuracy. Overall average accuracy, 96%. **(C)** Top: Large-field composite image of pooled cell populations from panel (A), displaying CFP2 (blue), mNG3Asp (green), and sfCherry3Csp (red) fluorescence channels. Scale bar: 500 µm. Middle: Population assignment map following GMM-based classification into 20 distinct categories. Bottom: Visualization of GMM classification confidence through membership probability mapping.

**Supplementary Figure S9.**
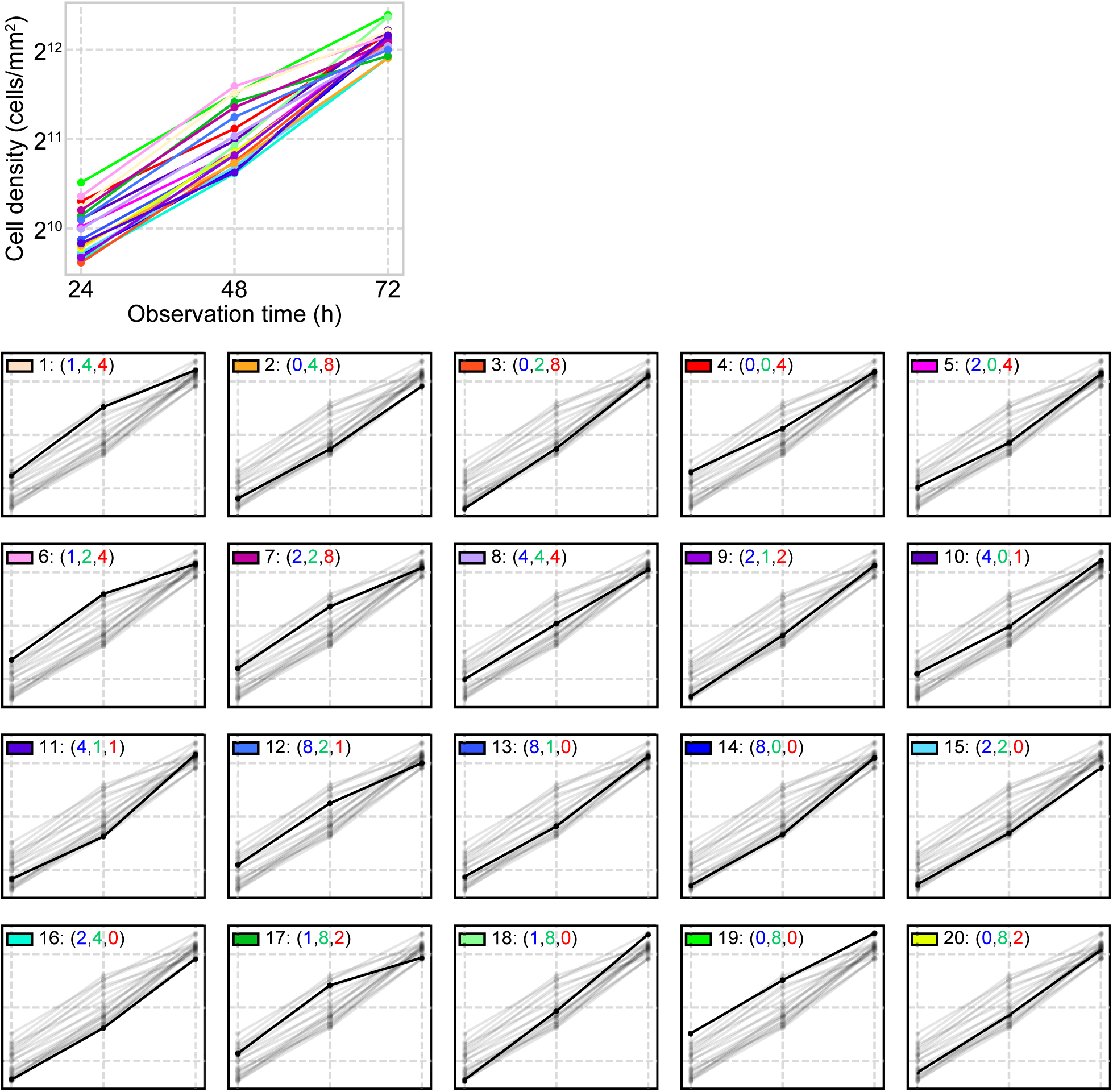
Cell growth of Caterpie labeled MDCK cells. Growth curves showing cell density of 20 different MDCK cell types fluorescently labeled with the Caterpie method measured every 24 hours. The lower panels display individual growth curves for each cell type.

